# Growth vs. Diversity: A Time-Evolution Analysis of the Chemical Space

**DOI:** 10.1101/2025.02.18.638937

**Authors:** Kenneth Lopez Perez, Edgar López-López, Flavie Soulage, Eloy Felix, José L. Medina-Franco, Ramon Alain Miranda-Quintana

## Abstract

Chemical space is a core and theoretical concept in cheminformatics, and it also has practical applications in drug discovery and other research areas. Chemical space is frequently associated with the number of molecules in the universe (e.g., chemical universe). It is well known that the number of compounds (both synthesized and theoretical ones) is rapidly increasing. It would be obvious to affirm that the chemical space is expanding (as a proxy of growth). But is the chemical diversity of compound libraries growing? In this study, we tackle this question by assessing quantitatively the time evolution of chemical libraries in terms of the chemical diversity as measured with molecular fingerprints. To tackle this task, we employed innovative cheminformatics methods to assess the progress over time of the chemical diversity of compound libraries available in the public domain. Using the iSIM and the BitBIRCH clustering algorithm, we conclude that, based on the fingerprints used to represent the chemical structures, just an increasing number of molecules cannot be directly translated to diversity for the analyzed libraries. With these tools, we have identified what releases contributed to the diversity of the library and the zones it did.

## 1. INTRODUCTION

Chemical space is a key concept in cheminformatics and molecular design.^1^ It serves as a systematic tool to analyze, study, and visualize the chemical diversity of all kinds of compounds contained in the “chemical universe”, which includes all compounds that can or could exist.^2,3^ The concept of chemical space has been defined and reviewed from different perspectives.^4,5^ For instance, Arús-Pous et al. describe it as “a concept to organize molecular diversity by postulating that different molecules occupy different regions of a mathematical space where the position of each molecule is defined by its properties".^6^ This definition has improved the development of new cheminformatic approaches with direct applicability in chemical diversity analysis, chemical structure classification, database design, virtual screening, and structure-property / structure-multiple property relationships. ^7,8^

Arguably there is not a unified definition of chemical space or chemical universe.^1^ The space definition can be constricted to certain regions of the “whole” space depending on the compounds included in it. For example, it has been estimated that the chemical space of small organic molecules exceeds 10^60^ compounds.^2,3^ It will also depend on the representations of the molecules, which can be chemical data^9^ (*e.g*., fingerprints^10^, physicochemical^11^ or quantum properties^12^), biological^13,14^ (*e.g*., bioactivity, bioavailability), or clinical^15^ (*e.g*., side effects) descriptors. This fact has brought up the idea of a consensus chemical space by combining multiple representations.^16^

Since the chemical space is too large to evaluate exhaustively^17^, the creation of field-specific libraries is important. Public repositories like ChEMBL^18^ contribute to the curation, standardization, organization, and filtration of chemical and biological information, and guarantee data accessibility to the public (*i.e*. academia and independent researchers)^19^, which is primordial in the current context of the accelerated accessibility to artificial intelligence, such as machine and deep learning methods.^20^

Parallelly, the accelerated increase in the number of compounds available in public repositories, and the creation of large (10^7^ compounds)^21^ and ultra-large (10^9^ compounds)^3,22^ libraries, opens new perspectives on the use and the study of the chemical universe. New strategies are required to navigate, cluster, and assess the diversity of these fast-increasing libraries; there is a necessity for more efficient models to do so.^17,23–25^

It is widely known that the cardinality in the explored chemical space is growing.^24,25^ However, it is also important to establish a rational quantification of its chemical diversity to have a clearer picture of its actual expansion. Also, providing information to guide the addition of new compounds in future releases. Just recently, Liu et al. reported a time-dependent comparison of different structural features of natural products obtained in a time series of the Dictionary of Natural Products, as compared to synthetic compounds^26^, which remarks on the importance of establishing systematic comparisons of the evolution of chemical diversity in chemical datasets.

In this contribution, we present several similarity-based tools to assess the time evolution of chemical libraries. The key component is the iSIM framework, which can efficiently quantify the intrinsic similarity (diversity) of each release thanks to its O*(N)* complexity. Moreover, the related notion of complementary similarity facilitates the study of how different zones of a library’s chemical space change over time. We also study if the diversity’s time evolution is dependent on the chosen fingerprint representation. Finally, the BitBIRCH clustering algorithm is used to dissect the evolving chemical spaces in a more “granular” way, looking at the formation of new clusters of compounds. The combination of these tools gives an unprecedented view on the formation of new chemical spaces, with varied resolutions, which can be broadly applicable to the design of novel compound libraries with specific desired functions. The adaptability and time efficiency make the proposed tools an attractive option for studying the chemical space of large molecular libraries.

## 2. METHODS

### 2.1 iSIM tools for dissecting chemical space

The similarity analysis of the different libraries was performed used iSIM as the basic tool. Traditional similarity indices are all based on the comparison between pairs of molecules, so they unequivocally scale as O(*N*^2^) when comparing *N* molecules. These steep computational cost is particularly pressing when dealing with millions of compounds, so given the expansive nature of commonly used libraries, we need other alternatives. iSIM bypasses the quadratic scaling problem by comparing all the molecules at the same time. The recipe for this is very simple: First, we arrange all the fingerprints in a matrix, and we add the elements of each of the columns, resulting in a vector *K* = [*k*_1_, *k*_2_, …, *k*_*M*_]. Clearly, *k*_i_ represents the number of “ones” in the *i*th column, which is all that we need to calculate the coincidence of “on” bits 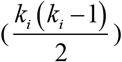, the coincidence of “off” bits 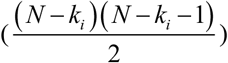, and the number of “on”-“off” clashes (*k*_*i*_ (*N* − *k*_*i*_)) in each column.

This information is enough to calculate the average value of multiple similarity indices. Here we focus on the Tanimoto similarity, T, as this well-known similarity index has been proven to correlate consistently in the ranking of compounds in structure-activity studies.^27,28^

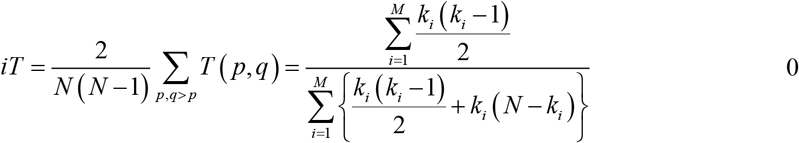

As indicated above, the iSIM Tanimoto (iT) value corresponds to the average of all the distinct pairwise Tanimoto comparisons, with the key advantage of being calculated in O(*N*). The iT thus corresponds to the internal diversity of the set (lower iT values indicate a more diverse collection of compounds). This is a global indicator of the diversity of the library, but it would be desirable to have local metrics for the evolution of chemical space. A simple way to do this is by utilizing the concept of complementary similarity. Since we can easily calculate the iT of a set, we can also easily calculate the set’s iT after a molecule has been removed from it. This is known as the complementary similarity of the removed molecule. Lower complementary similarity values correspond to molecules that are central to the library (they are medoid-like), and high complementary similarity values are indicative of outlier molecules, in the periphery of the set. After identifying the central and outlier regions of the set, we can then calculate their corresponding iSIM values and see how their internal diversity changes over time. Additionally, we compared the medoids and outliers between releases of the set by calculating the iSIM when merging them. To shed more light on the time of evolution of different sectors of chemical space we analyzed how different iterations of a given library are related to each other. For this, we used the set Jaccard similarity index, J:

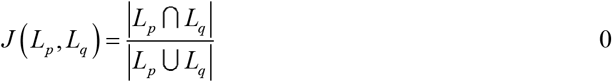

where *L*_*p*_ and *L*_*q*_ represent a sector (medoid or outliers) of a library, but correspond to years *p* and *q*. Where for a given set X, |X| indicates the cardinality (size) of the set, for Eq. 2 the cardinality of the intersection over the cardinality union of the sectors of the libraries. We define the medoids as molecules in the lowest 5th percentile according to complementary similarity values, and the outliers as the ones in the highest 5th percentile.

Finally, to gain even more insights into the inner structure of the chemical space, we recourse to clustering. Once again, the O(*N*^2^) scaling of standard clustering techniques, like Taylor-Butina^29^ and Jarvis-Patrick^30^, severely limits the scope of the chemical space that could be explored. For this reason, we decided to use the recently proposed BitBIRCH^31^ algorithm. This method draws inspiration from the Balanced Iterative Reducing and Clustering using Hierarchies (BIRCH)^32^ by using a tree structure to reduce the number of comparisons required to group all the data. The key difference is that the original BIRCH algorithm could only be used to cluster objects represented with continuous vectors, and using the Euclidean distance. BitBIRCH, on the other hand, relies on iSIM to process binary vectors and to use the Tanimoto similarity.^31^

### 2.2 Chemical libraries

#### ChEMBL

ChEMBL^33^ is a large-scale, high-quality, manually curated, open Global Core Biodata Resource focused on drug-like bioactive compounds. The data are manually extracted from primary scientific literature, updated regularly, and rigorously curated and standardized. In addition to scientific literature data, ChEMBL incorporates deposited datasets, including those from neglected disease screening programs, crop protection research, drug metabolism and disposition studies, bioactivity data extracted from patents, and chemical probe screening programs. Currently, ChEMBL contains over 20 million bioactivity measurements for more than 2.4 million compounds and 15,500 protein targets. We analyzed the complete versions of ChEMBL from releases 1 through 33. In addition, we applied our framework only to the natural products of each of the releases.

#### DrugBank

DrugBank^34,35^ is an online database containing information on drugs and drug targets, making it highly used in cheminformatics and bioinformatics. For targets, it includes sequences, structures, and pathways. It combines chemical, pharmacological, and pharmaceutical data for drugs. We analyzed the releases of drugs included in the database from 2005 to 2022.

#### PubChem

PubChem^36^ is a widely used open chemistry database put to disposition by the National Institute of Health (NIH). It was launched in 2004, since it has added information on millions of small compounds and larger molecules. PubChem compiles information from several sources including government, agencies, publications, and patents. For each compound, PubChem contains multiple identifiers, physical and chemical properties, toxicological information, related compounds, etc. For this publication, we analyzed the releases from 2004 to 2023.

### 2.3. Molecular representations

It is well known that molecular similarity measurements depend on the representations^37^, thus we decided to carry out this study with three different types of 2048-bit fingerprints: RDKit^38^, ECFP4^39,40^ (radius = 2), and ECFP6^39,40^ (radius = 3). All fingerprints were calculated using the FPSim2^41^ python module using SMILES strings obtained from the above-mentioned libraries.

## 3. RESULTS AND DISCUSSION

### 3.1 Absolute Size & Diversity

We will illustrate the key points of our analysis using the ChEMBL library, however, the same conclusions can be observed on other libraries presented in the Supplementary Information. Overall, a simple counting strategy is not enough to evaluate the potential expansion of chemical space. As shown in Fig. 1 a), ChEMBL (as virtually every library) always increases its number of compounds over time, but this is not necessarily a direct proxy of the chemical diversity of the set, nor an obvious connection to new chemistries being explored. As stated in the Introduction, in this work we did not consider the chemical space expansion as a direct synonym of the increasing number of compounds in chemical libraries. Figure 1 b) shows how the average Tanimoto similarity of the whole library (we will name as iSIM for the rest of the text) does not fluctuate much with the increase of the library’s size. We want to remark that this calculation is extremely easy to calculate over time, as we only need to add the column-wise sum of the new incoming molecules from release to release to calculate the new iSIM value.

**Figure 1:**
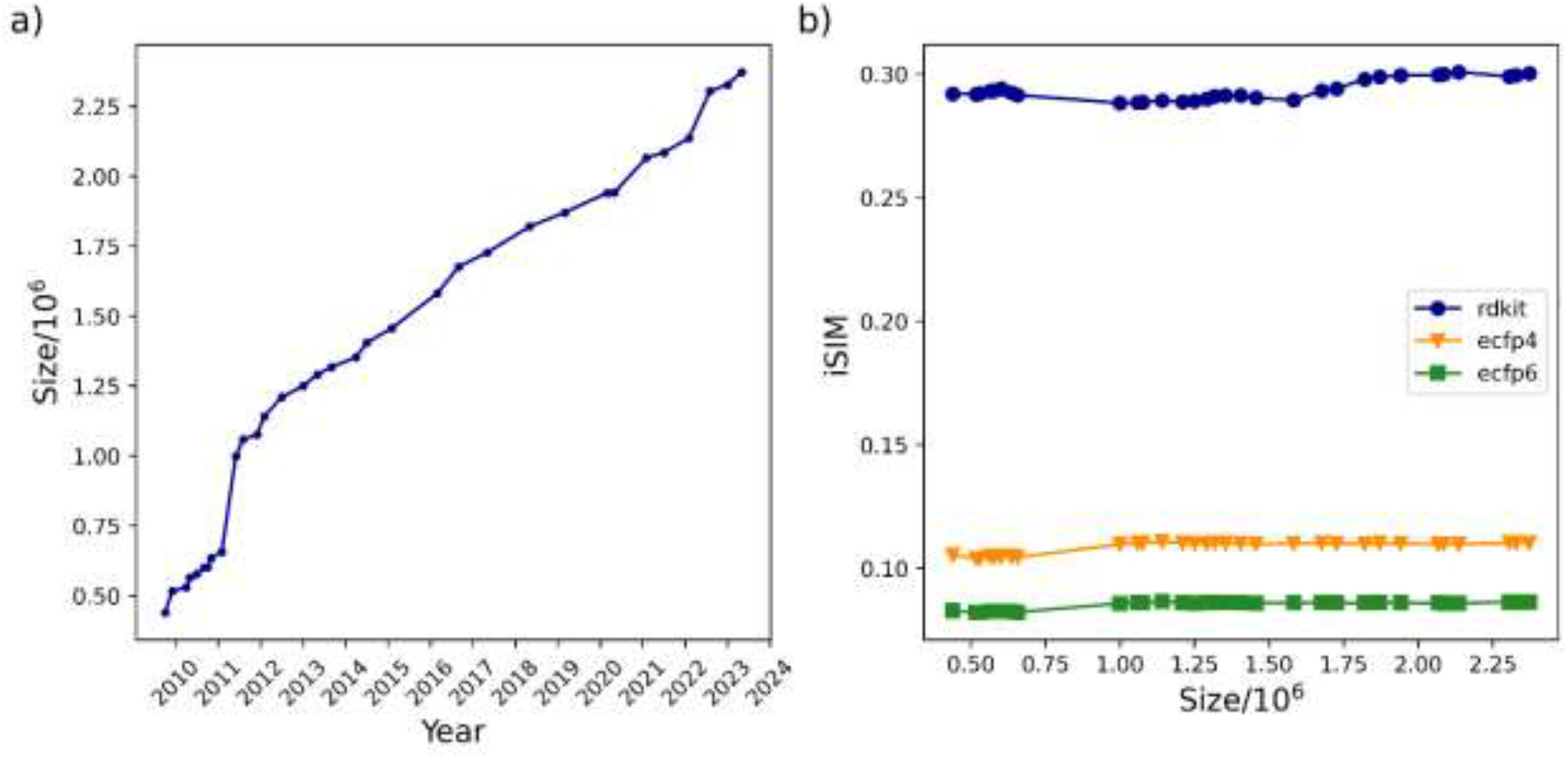
(a) Size evolution of the ChEMBL database across releases (1-33) over time and (b) the variation of iSIM-Tanimoto with database size for the same ChEMBL releases.

The same trend in Fig. 1 b) can also be seen in Fig. 2, iSIM of the whole library, as well as the central and outlier regions remained essentially constant since 2011, despite more than 1 million molecules being added since then. As expected, the iSIM value of the whole set is somewhere in between that of the medoids (which are notably less diverse) and the outliers (which are markedly more diverse). It seems like, after an initial period of small fluctuations before 2011, the average similarity of the whole set and the medoids essentially reaches an “equilibrium”. In the case of the outliers, we see the most notable changes in the iSIM value before 2015. The same steady trends in the iSIM value over the ChEMBL releases are observed with other similarity indexes like Russell-Rao and Sokal-Michener. (SI: Fig. S1) When focusing on ChEMBL’s natural products there is a noticeable decrease in diversity from the first to the second release. (SI: Fig S10) In the case of the natural product’s outliers, it is also clear how in 2011 and 2015 the additions contributed to increasing the diversity of the set. (SI: Fig S9) PubChem shows a different trend in the iSIM value over time; during the first 10 years, the diversity increases (similarity decreases) overall, after the 2016 addition the behavior changed, the iSIM values have been slowly increasing since. (SI: Fig 26)

**Figure 2:**
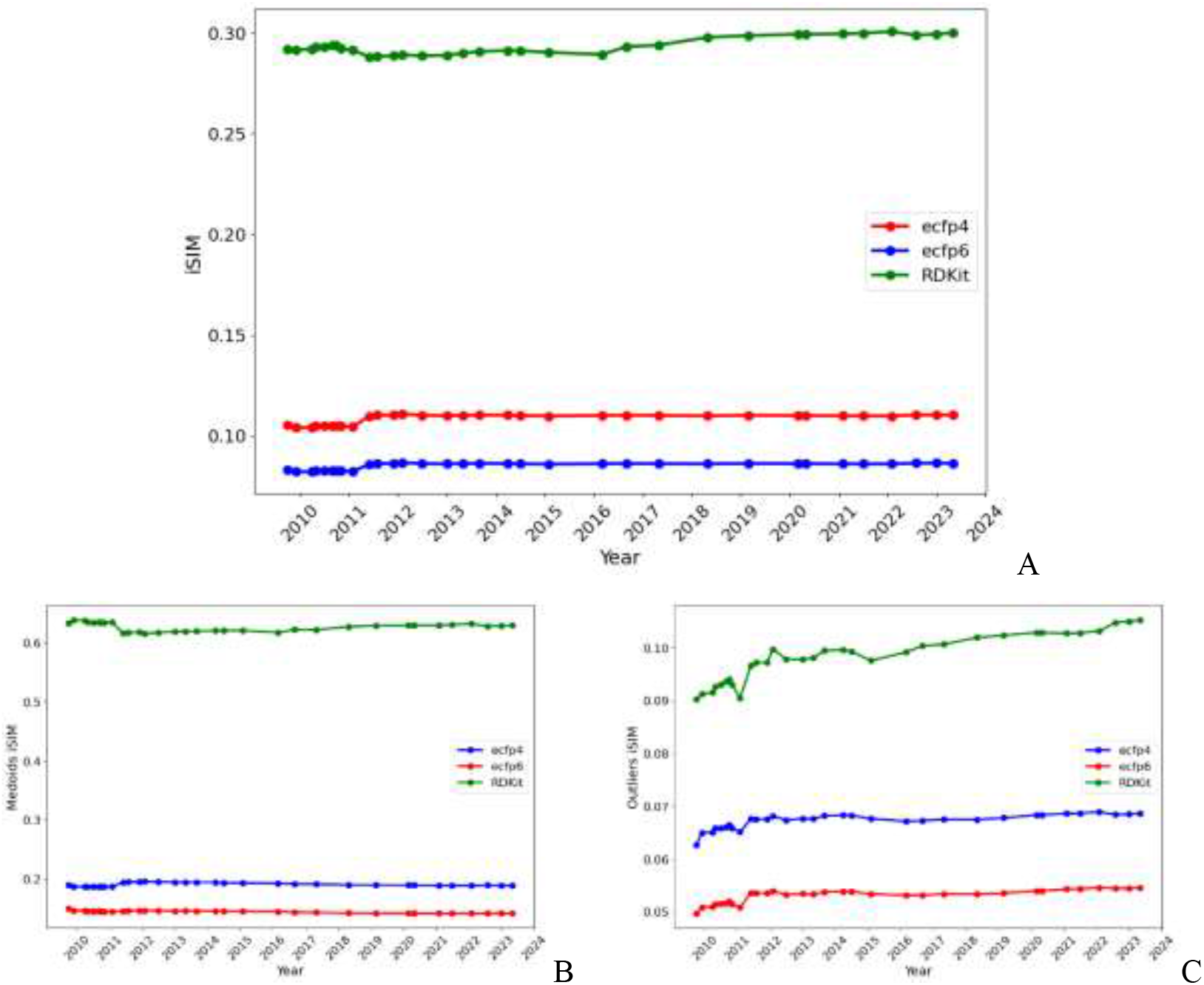
Variation of iSIM over time for the (A) entire ChEMBL database, (B) medoids, and (C) outliers. Medoids and outliers are considered the 5% of the set with the lowest/highest complementary similarity, respectively.

### 3.2 iSIM Speed

An interesting exercise is to check the actual rate of change of the iSIM values over time, which we termed “iSIM speed”. In Fig. 3 we show that the rate of change over time is relatively similar for the whole set and for the medoids and outliers. However, the type of fingerprint used to represent the molecules has a great impact even in the qualitative direction of the change. For example, in the whole set and medoid region the RDKit fingerprints reflect a decrease in iSIM, while the ECFP fingerprints show the opposite change. That is, the RDKit fingerprints suggest an increase in diversity, while ECFP representation shows that the new molecules actually decreased the overall separation between the compounds in the library. It is also notable that with passing years the ECFP fingerprints tend to show a more stable landscape, while RDKit fingerprints show comparatively bigger fluctuations. As for the outliers, the changes are more consistent between the three molecular representations. Overall, the greatest change in iSIM was in 2011 as appreciated in the three panels of Fig. 3; this date matches one of the biggest increases in the size of the ChEMBL library (Fig. 1a). These changes are also appreciated in the analogous iSIM speed respect to the size (d[iSIM]/d[size]) shown in the SI (Fig. S2).

**Figure 3:**
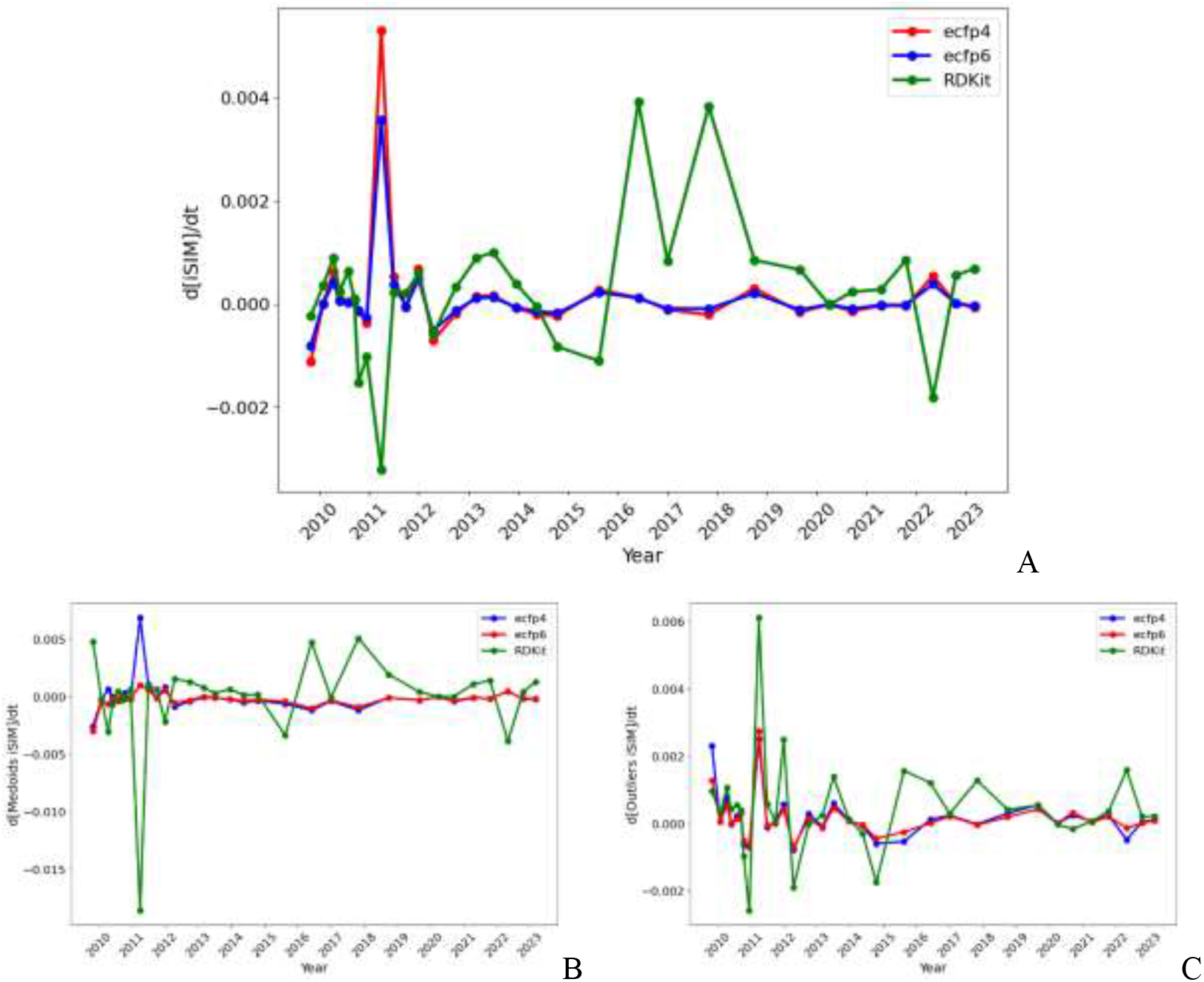
iSIM speed with respect to time for the A) entire, B) the medoids, and C) outliers of the ChEMBL library.

### 3.3 Jaccard set similarity for medoids and outliers

Fig. 4 shows the Jaccard set similarity between releases for medoids and outliers in the ChEMBL library. It suggests that in virtually every iteration there is a strong overlap between the core of the library and, perhaps more surprisingly, even between the outliers. Notice that the behavior of the medoids and outliers is strikingly similar, both showing the same overall trends, even while corresponding to drastically different regions of chemical space. It is interesting that in this representation it is easier to see how there are some “pivotal” updates in the datasets that are preserved throughout multiple years. The main observed change is in 2011, which matches with the observations from the previous section. It is also apparent how the latest updates seem to be highly correlated with each other, in particular since 2019-2020. These trends are observed also with the ECFP4 and ECFP6 fingerprints (SI: Fig. 3-4).

To complement the results shown in Fig. 4, we calculated the iSIM value of the merged zones of chemical space between releases. The same observations were obtained (Fig. S5), the main change in the iSIM value happened in 2011 for both medoids and outliers. Structures of the outlier (molecule with the largest complementary similarity) and medoids (molecule with the lowest complementary similarity) from each year are included in the SI.

**Figure 4:**
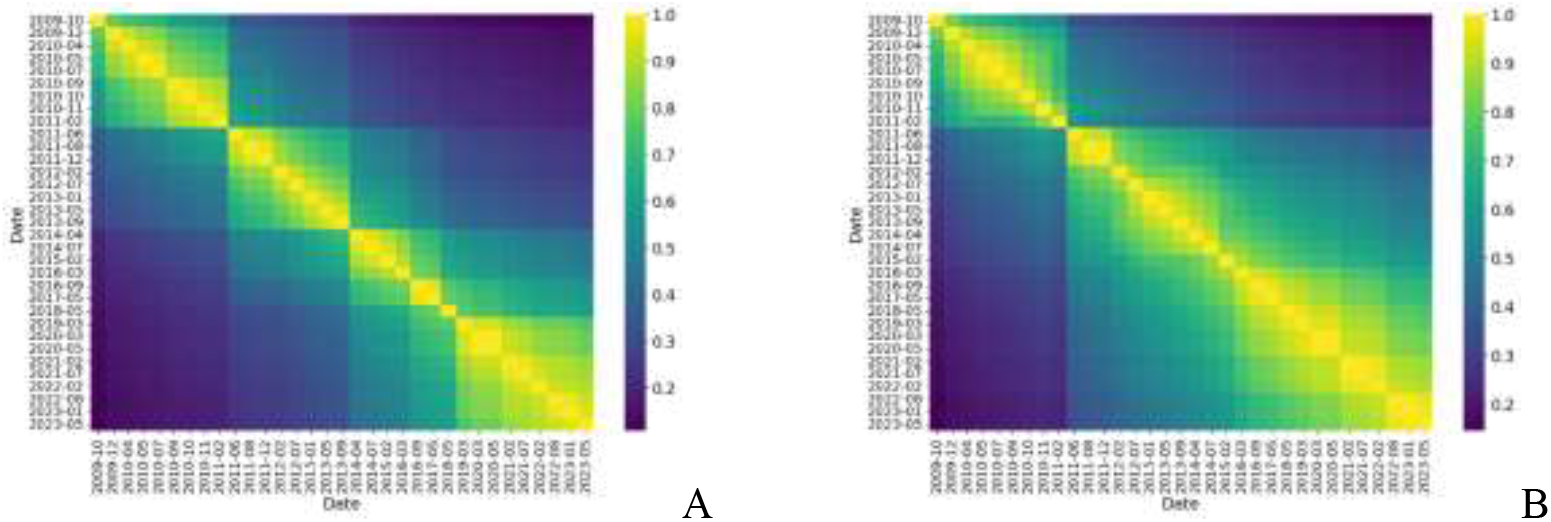
Jaccard set similarity values of the medoid (A) and outlier (B) regions of the ChEMBL library represented with RDKit fingerprints.

### 3.4 Clustering

Finally, we studied how the BitBIRCH clustering of ChEMBL has evolved through time, we performed it with a threshold of 0.65, as this value is in the relevant range for drug design and has shown stability in previous works. We define dense clusters as the ones with more than 10 elements, and outliers as the ones with less. First, notice how the average similarity of the top 10 most populated clusters shows an overall decreasing tendency (Fig. 5A), while their average population is increasing (Fig. 5B). This is expected of these sphere exclusion-like clustering algorithms, which tend to produce slightly more diffuse clusters with increasing number of members. However, perhaps more telling towards the actual library expansion, both the number of dense and outliers clusters (Fig. 5C-D) show a decisively upward tendency, indicating that the incoming molecules with each iteration are not only going to over-explored regions of chemical space but are also covering new regions in the library’s scope. With extended connectivity fingerprints (ECFP4 and ECFP6) the populations and number of dense/outlier clusters have the same increasing tendency as the RDKit fingerprints. However, the average iSIM of most populations has more fluctuations and does not have a decreasing tendency (SI: Fig. S6-S7).

**Figure 5:**
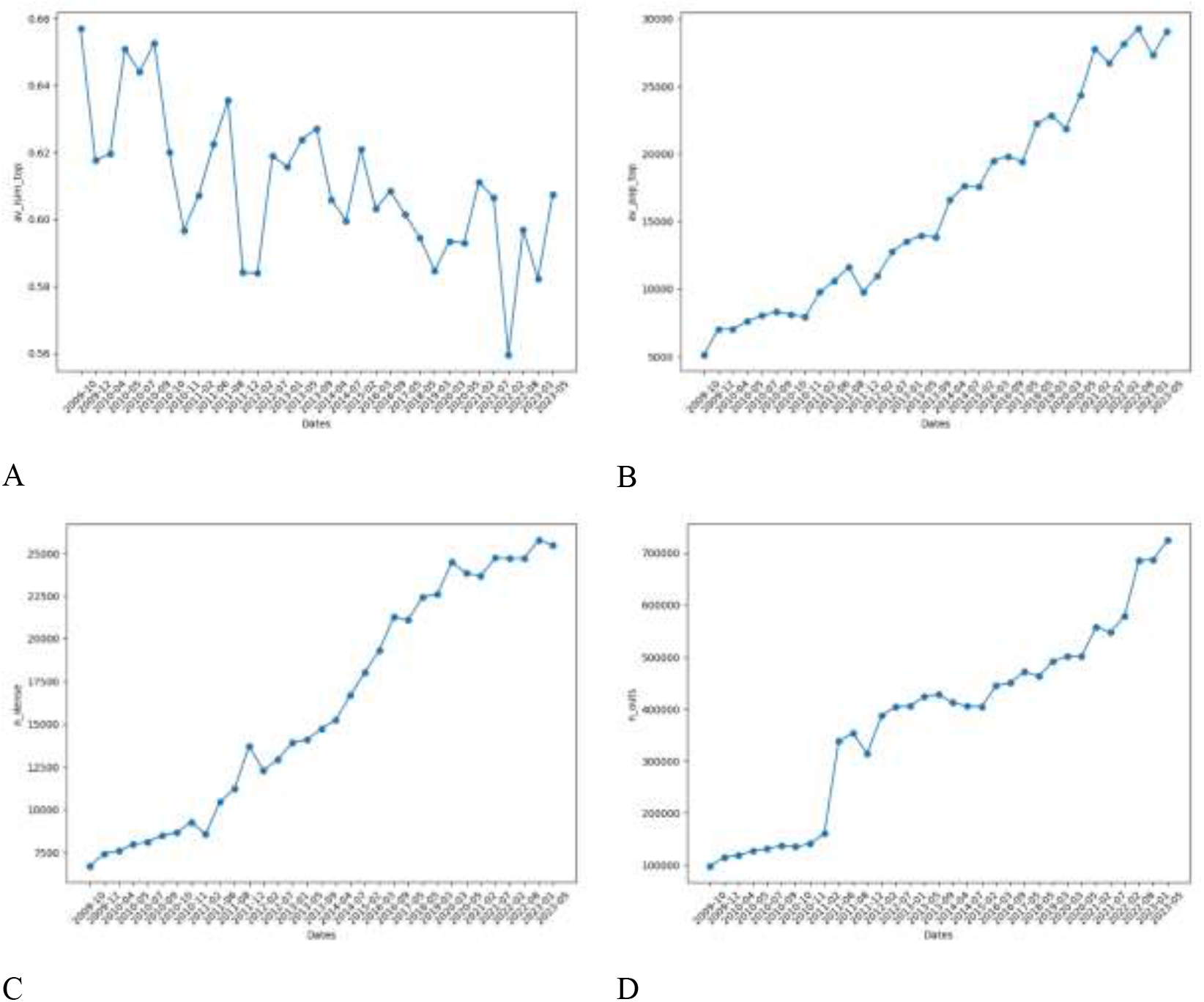
(A) Average iSIM of the top 10 most populated clusters (B) Average population of the top 10 most populated clusters (C) Number of dense clusters (D) Number of outliers for the BitBIRCH clustering of the ChEMBL releases over time represented with RDKit fingerprints.

## 4. CONCLUSIONS AND OUTLOOK

In this work, we have shown how increasing the number of molecules in a molecular library cannot be directly translated to more diversity or “expansion” of the chemical space. The average similarity of the libraries, iSIM Tanimoto, tends to be a rather stable quantity, not reflecting the impact of the expansion in chemical space (considering the compounds deposited in the public databases studied in this work: heavily focused on drug discovery projects). Different molecular representations can lead to different conclusions regarding the increase or not of chemical diversity in abrupt changes in the chemical space. However, the overall tendencies over time do not vary much across the types of fingerprints studied, they agree on which years there are significant changes in the diversity. Across the multiple releases of the ChEMBL database, the medoid and outlier regions tend to be conserved over multiple years, according to their Jaccard similarity the only significant observed change was in 2011.

The clustering analysis provides a more nuanced view of the expansion of chemical space and the clearest indication of an actual expansion, as reflected in the number of newly generated dense clusters and outliers.

It remains to be explored quantitatively if the continued enumerated chemical libraries (large and ultra-large libraries) are increasing the chemical diversity of the chemical space or just increasing the number of molecules (as it happened in the 1990s with the “traditional” combinatorial libraries). Due to the flexibility and adaptability of the proposed framework, it can be used to evaluate the chemical diversity of the libraries based on other types of molecular representations such as chemical scaffolds, and continuous properties (drug-like, ADMETox, constitutional descriptors, etc.) in future work. It remains to evaluate the chemical diversity of the specialized libraries based on other types of molecules such as small molecules, peptides, macrocycles, metallodrugs, etc.

## ACKNOWLEDGEMENTS

KLP, FL, and RAMQ thank the National Institute of General Medical Sciences of the National Institutes of Health for support under award number R35GM150620. E.L.-L. is grateful to Consejo Nacional de Humanidades, Ciencia y Tecnología (CONAHCyT), Mexico, for the Ph.D. scholarship number, 894234. E. L.-L. also thanks DrugBank for providing academic access to their platform, which was used to explore the data of the approved drugs presented in this work. We also thank the funding of DGAPA, UNAM, Programa de Apoyo a Proyectos de Investigación e Innovación Tecnológica (PAPIIT), grant No. IG200124.

## DATA AND AVAILABILITY

### Datasets

The SMILES strings were obtained from the publicly available websites:

ChEMBL^18^ https://chembl.gitbook.io/chembl-interface-documentation/downloads

DrugBank^34^ https://go.drugbank.com/releases/latest

PubChem^36^ https://pubchem.ncbi.nlm.nih.gov/docs/downloads

### Code and análisis

iSIM functionalities, BIRCH clustering algorithm, and examples of how to use them for the diversity vs growth analysis of ChEMBL’s natural products are included in the linked repository. https://github.com/mqcomplab/ChemUniverse

