## Supplementary Information for "Growth vs. Diversity: A Time-Evolution Analysis of the Chemical Space"

4.

- *ChEMBL*

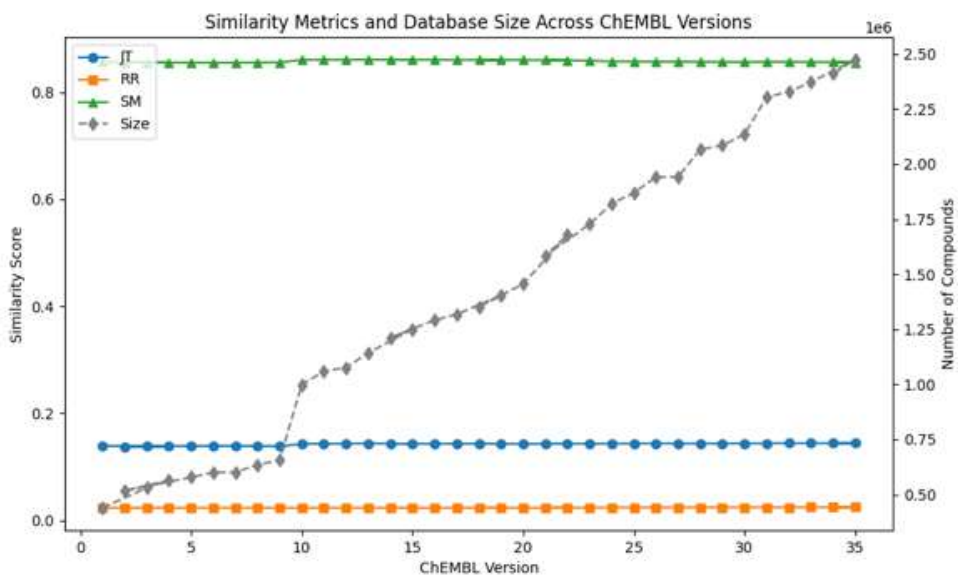

**Figure S1.** Variation of iSIM over time for the entire ChEMBL database with three different similarity indexes Jaccard-Tanimoto (JT), Russell-Rao (RR), and Sokal-Michener.

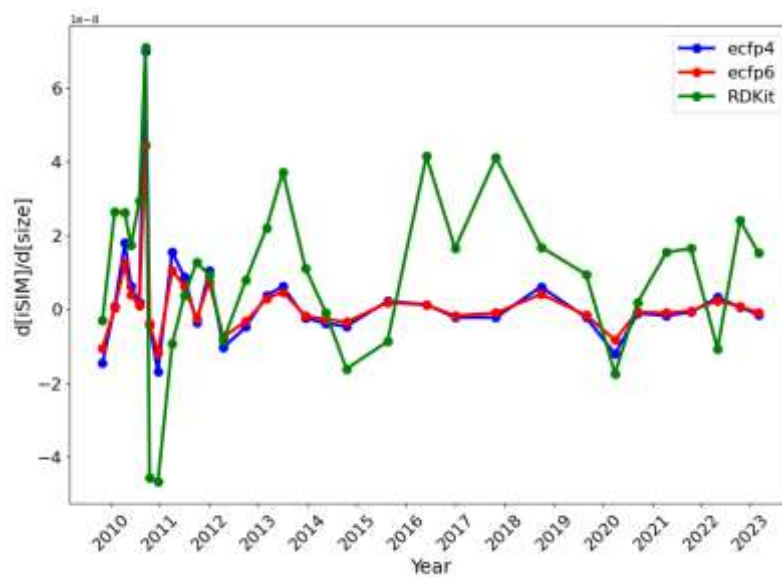

A

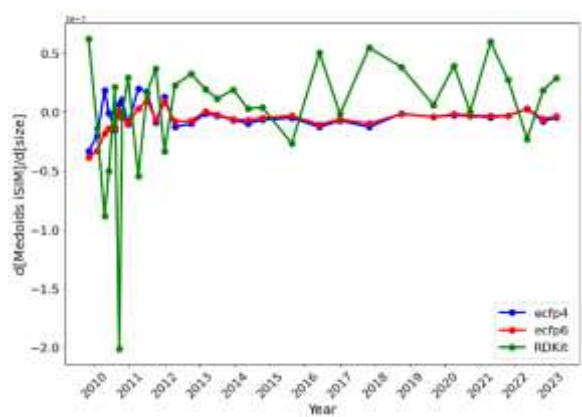

B

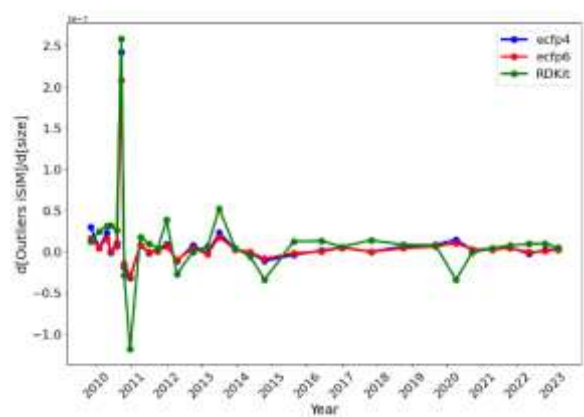

C

**Figure S2.** iSIM speed respect to size for the A) entire, B) the medoids, and C) outliers of the ChEMBL library.

A

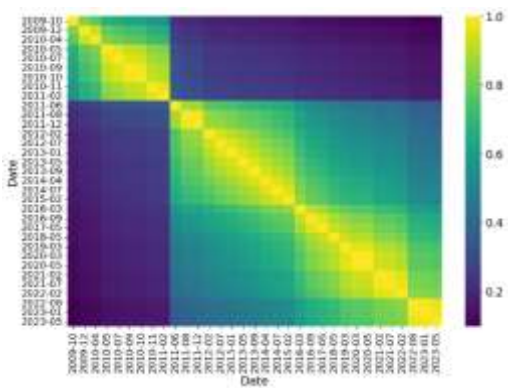

B

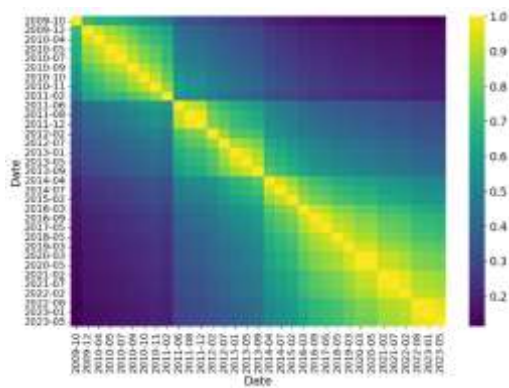

**Figure S3.** Jaccard set similarity values of the medoid (A) and outlier (B) regions of the ChEMBL library releases represented with EFCP4 fingerprints.

A

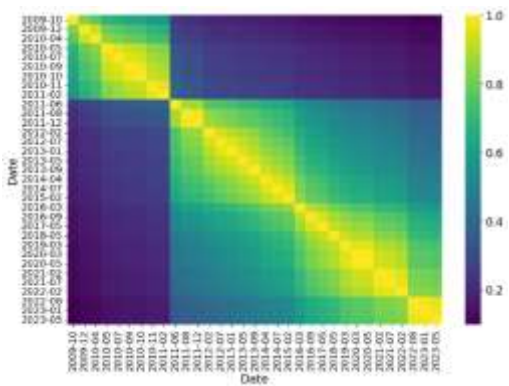

B

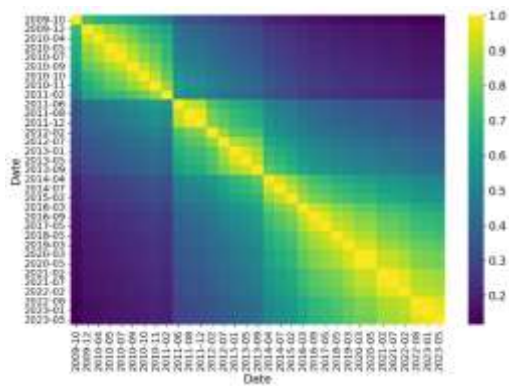

**Figure S4.** Jaccard set similarity values of the medoid (A) and outlier (B) regions of the ChEMBL library releases represented with EFCP6 fingerprints.

A

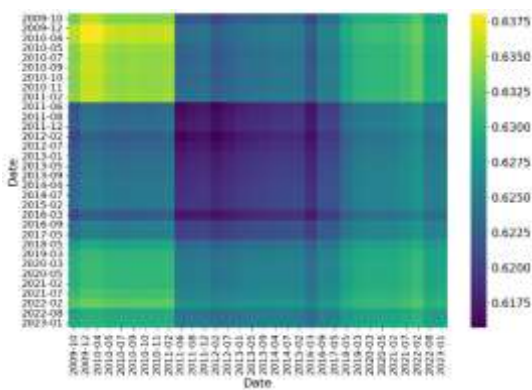

B

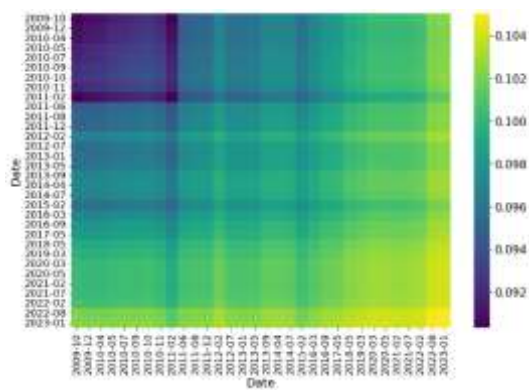

**Figure S5.** iSIM Tanimoto of the merged medoid (A) and outlier (B) regions of the ChEMBL library releases represented with RDKit fingerprints.

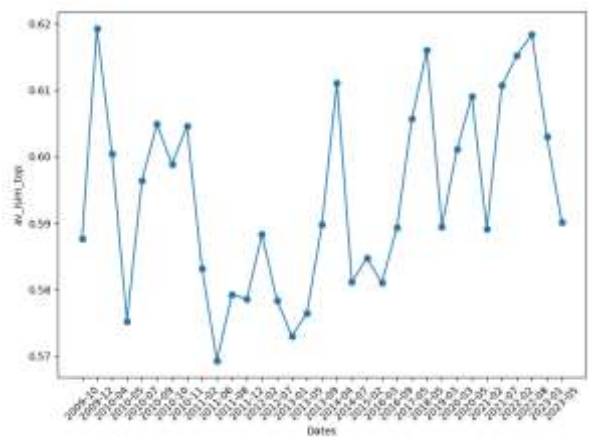

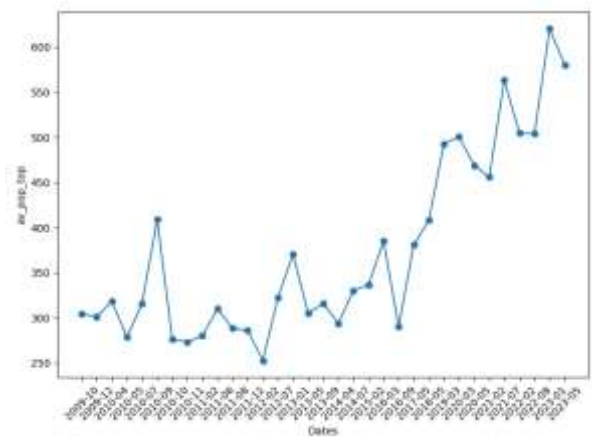

A

B

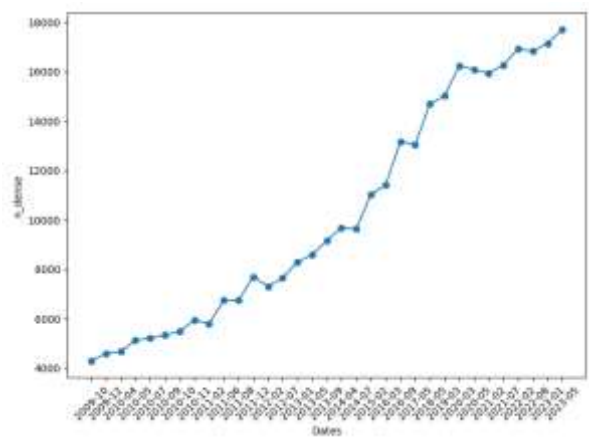

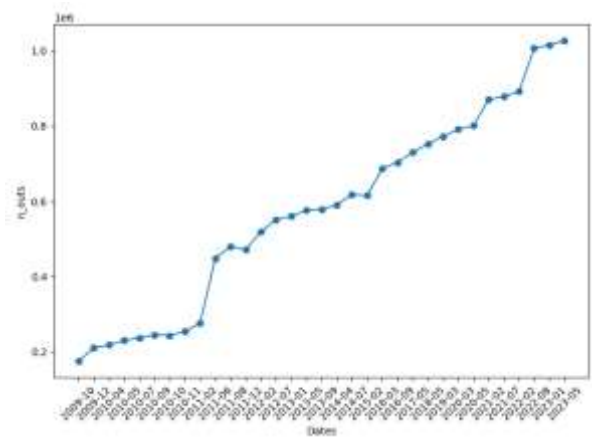

C

D

**Figure S6.** (A) Average iSIM of the top 10 most populated clusters (B) Average population of the top 10 most populated clusters (C) Number of dense clusters (D) Number of outliers for the BitBIRCH clustering of the ChEMBL releases over time represented with ECFP4 fingerprints.

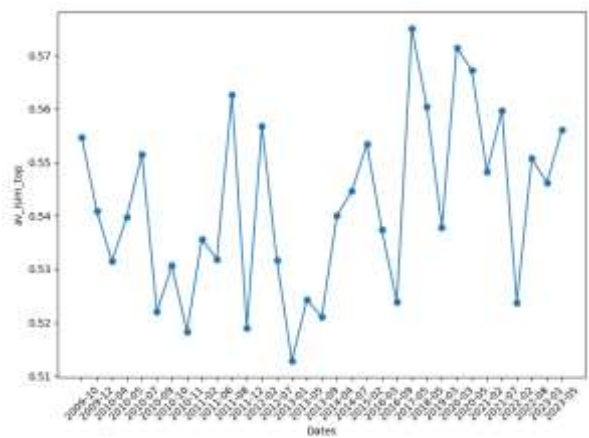

A

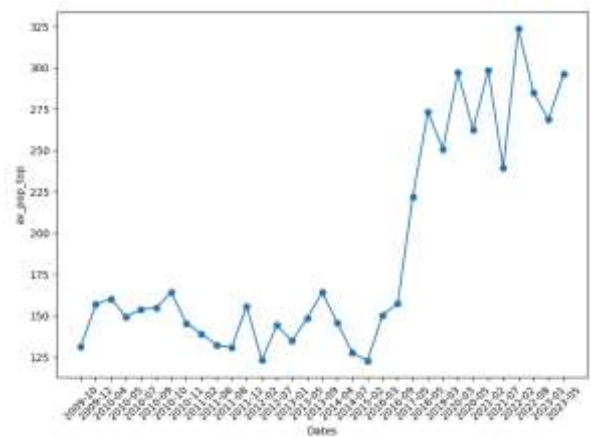

B

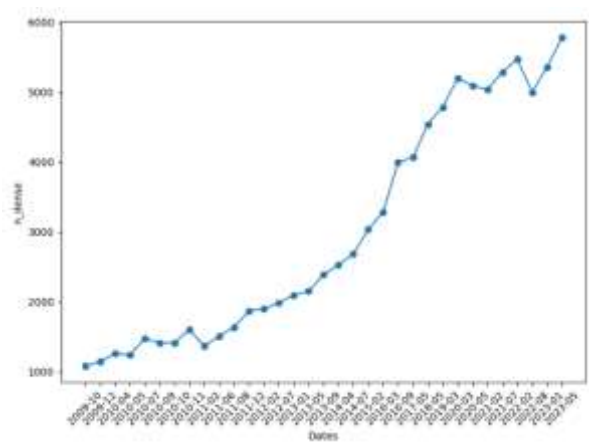

C

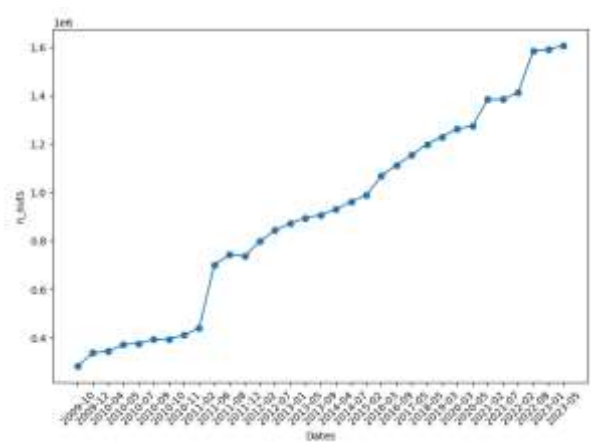

D

**Figure S7.** (A) Average iSIM of the top 10 most populated clusters (B) Average population of the top 10 most populated clusters (C) Number of dense clusters (D) Number of outliers for the BitBIRCH clustering of the ChEMBL releases over time represented with ECFP6 fingerprints.

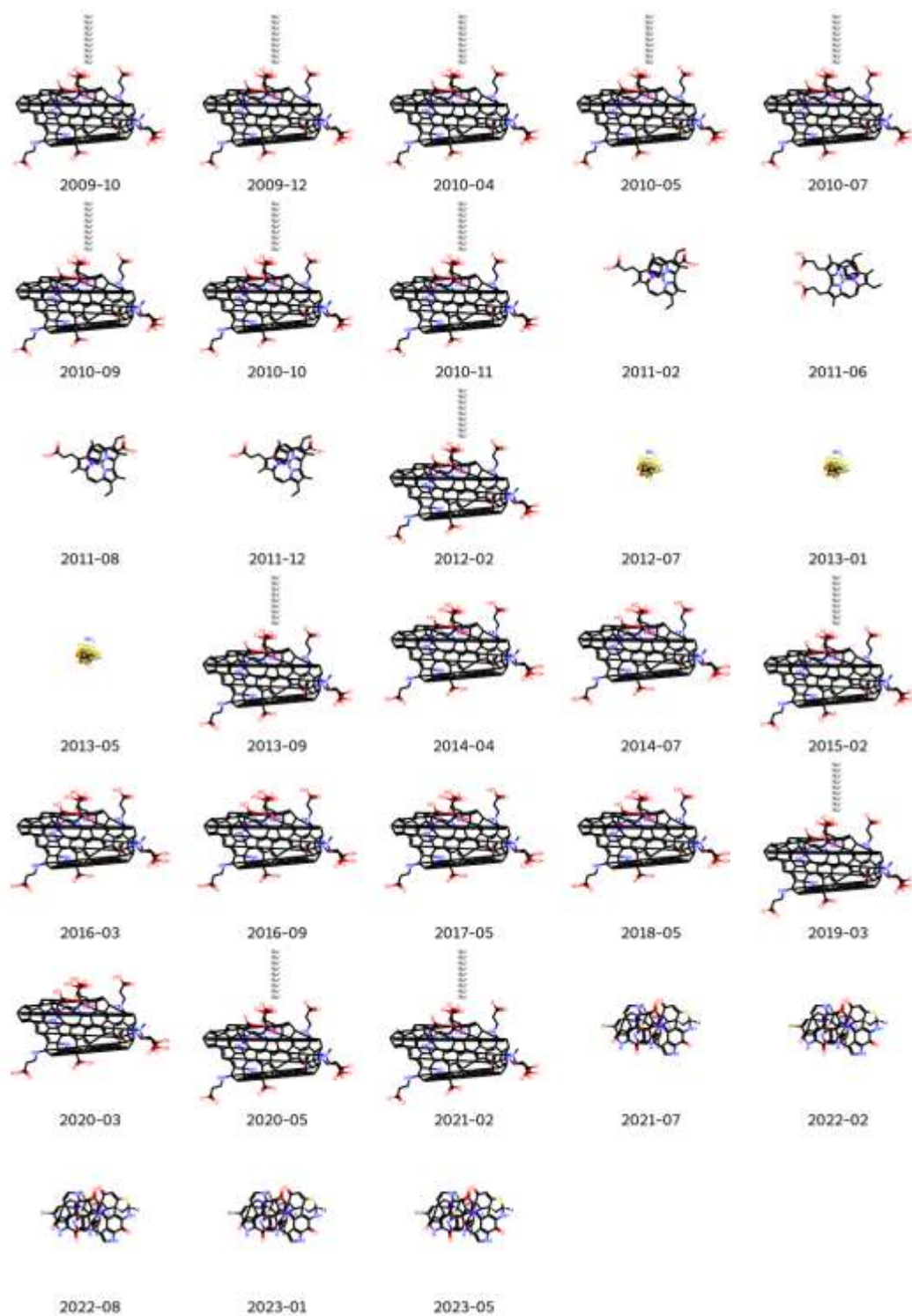

**Figure S8.** Medoids over time for ChEMBL.

|  |  |  |  |  |
| --- | --- | --- | --- | --- |
| $^{123}\text{I}^-$ | $\text{I}^-$ | $\text{Cl}^-$ | $\text{Co}^{2+}$ | $\text{F}^-$ |
| 2009-10 | 2009-12 | 2010-04 | 2010-05 | 2010-07 |
| $\text{Li}^+ \text{Cl}^-$ | $\text{H}_2\text{O}$ | $\text{Mn}^{2+}$<br>$\text{Cl}^- \text{Cl}^-$ | $\text{Os}^{3+}$ | $\text{Na}^+$ |
| 2010-09 | 2010-10 | 2010-11 | 2011-02 | 2011-06 |
| $\text{Hg}^{2+}$<br>$\text{Cl}^- \text{Cl}^-$ | $\text{NH}_4^+$ | $\text{H}_2\text{S}$ | $\text{O}^{2-} \text{Cu}^{2+}$ | $\text{Br}^-$<br>$\text{CaBr}^-$ |
| 2011-08 | 2011-12 | 2012-02 | 2012-07 | 2013-01 |
| $\text{Ba}^{2+}$<br>$\text{HOHO}^-$ | $\text{NaCl}^-$ | $\text{Ba}^{2+}$<br>$\text{HOHO}^-$ | $\text{Cl}^-$<br>$\text{Mg}^{2+}$ | $\text{Ba}^{2+}$<br>$\text{Cl}^-$<br>$\text{H}_2\text{CH}_2\text{O} \text{Cl}^-$ |
| 2013-05 | 2013-09 | 2014-04 | 2014-07 | 2015-02 |
| $^{133}\text{Xe}$ | $^{133}\text{Xe}$ | $\text{SbH}_3$<br>$\text{H}_2\text{S}$<br>$\text{H}_2\text{S}$<br>$\text{H}_2\text{SH}_2\text{SH}_2\text{S}$ $\text{SbH}_3$ | $\text{Cl}$ | $^{123}\text{I}^-$ |
| 2016-03 | 2016-09 | 2017-05 | 2018-05 | 2019-03 |
| $\text{H}_2\text{Se}$ | $^{223}\text{Ra}$ | $\text{H}^{76}\text{Br}$ | $^{223}\text{Ra}$ | $^{223}\text{Ra}$ |
| 2020-03 | 2020-05 | 2021-02 | 2021-07 | 2022-02 |
| $^{82}\text{Rb}$ | $\text{Na}^{124}\text{I}^-$ | $^{131}\text{Cs}$ | | |
| 2022-08 | 2023-01 | 2023-05 |  |  |

**Figure S9.** Outliers over time for ChEMBL.

- *ChEMBL natural products*

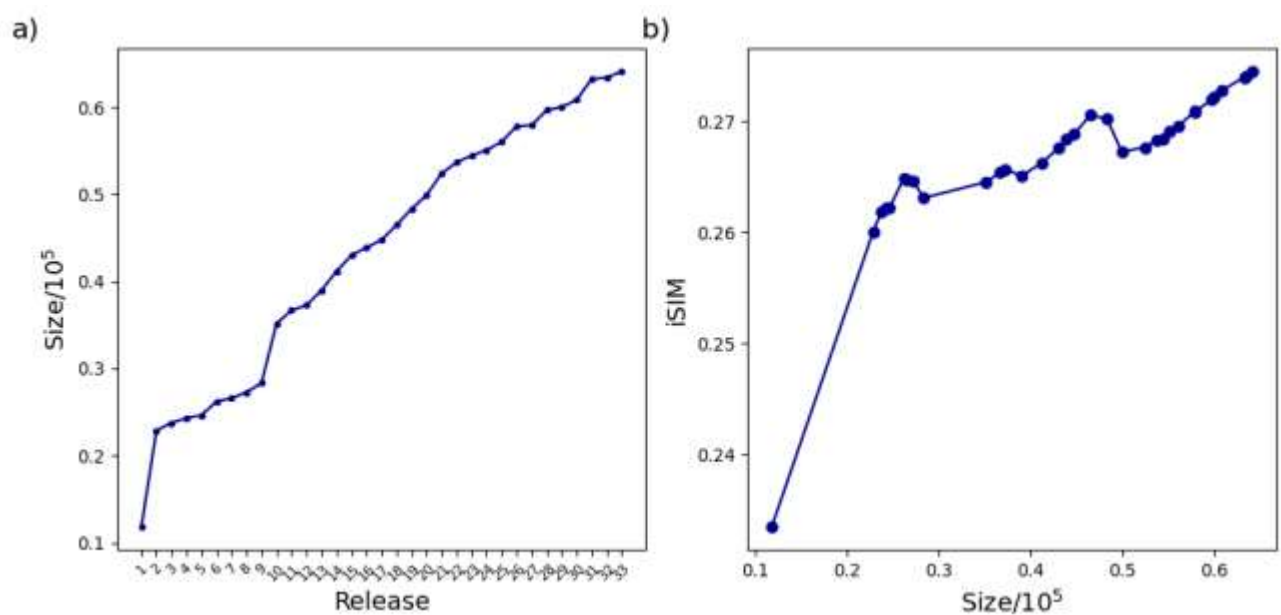

**Figure S10.** (a) Size evolution of the ChEMBL natural products across releases (1-33) over time and (b) the variation of iSIM-Tanimoto with database size for the same ChEMBL natural products releases represented with RDKit fingerprints.

A)

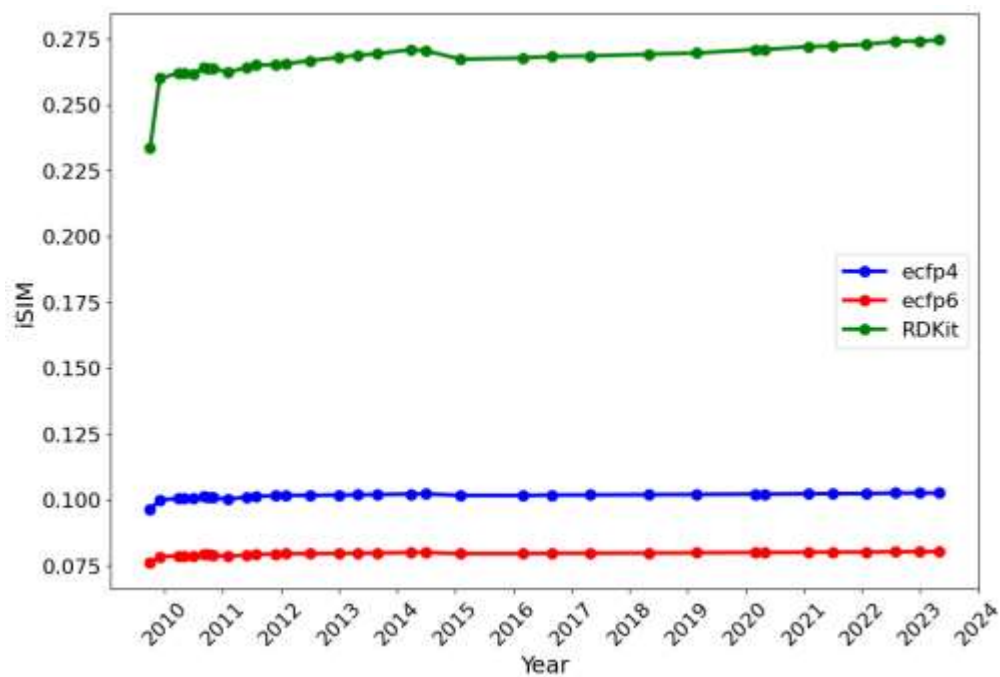

B)

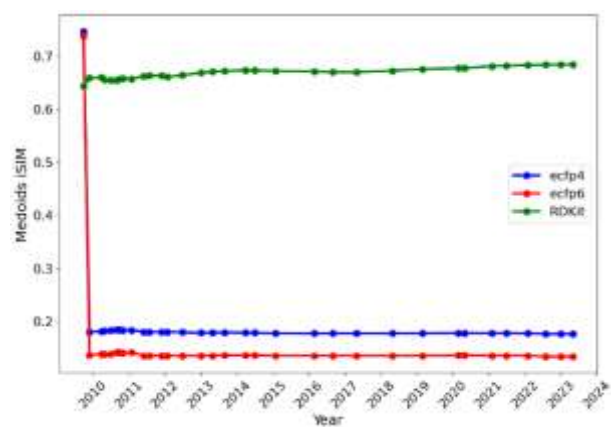

C)

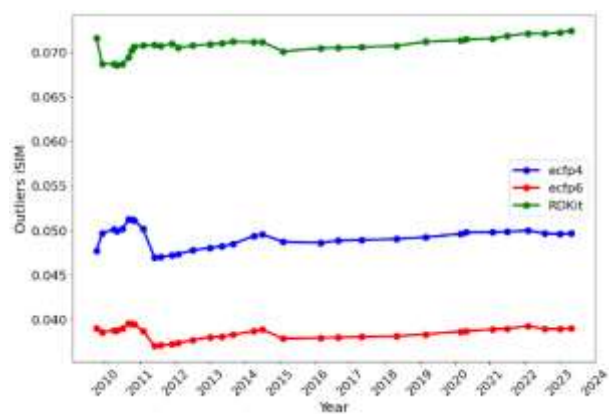

**Figure S11.** Variation of iSIM over time for the (A) ChEMBL natural products, (B) medoids, and (C) outliers. Medoids and outliers are considered the 5% of the set with the lowest/highest complementary similarity, respectively.

A)

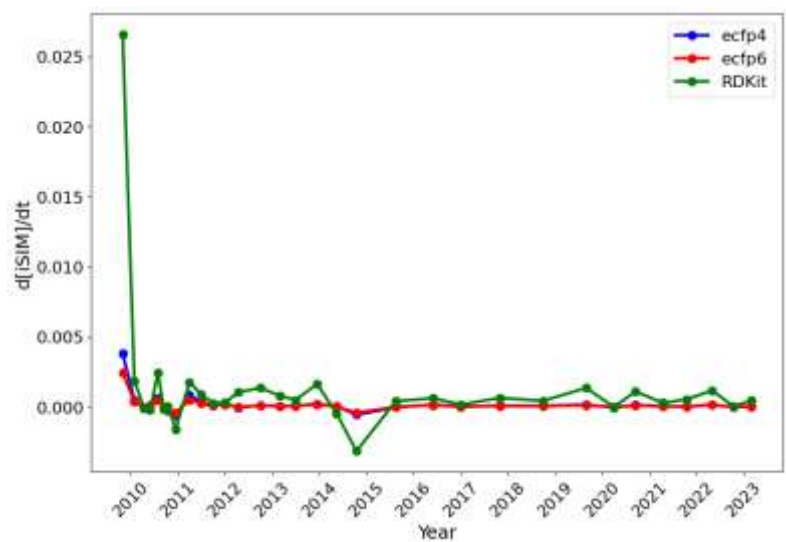

B)

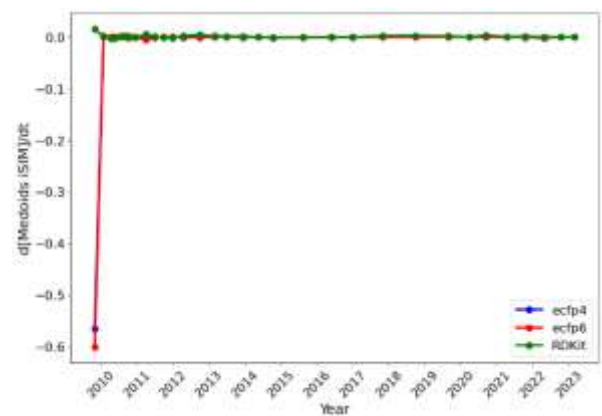

C)

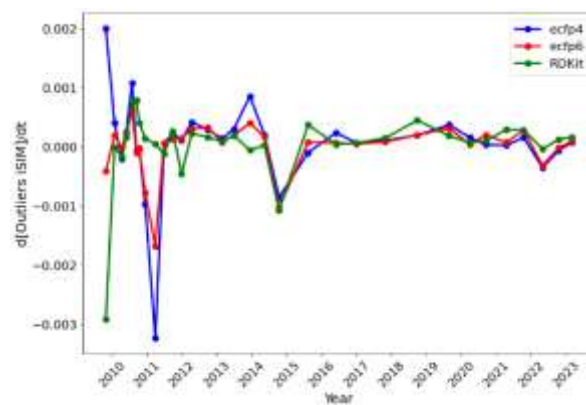

**Figure S12.** Variation of iSIM over time for the (A) entire ChEMBL natural products, (B) medoids, and (C) outliers. Medoids and outliers are considered the 5% of the set with the lowest/highest complementary similarity, respectively.

A)

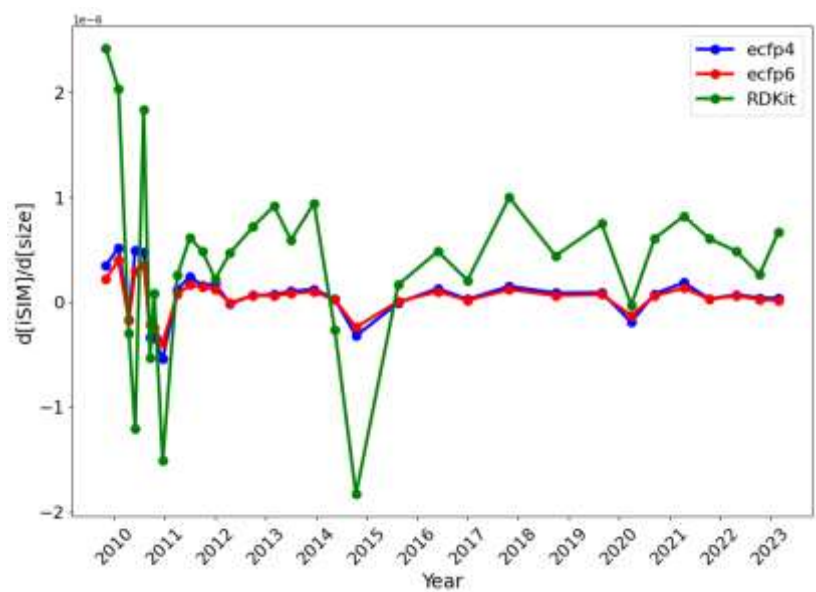

B)

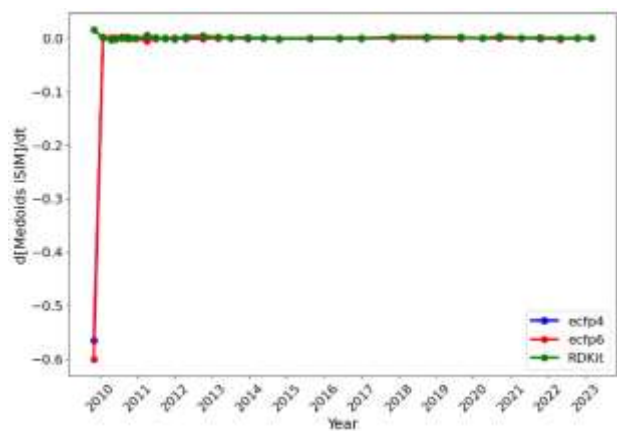

C)

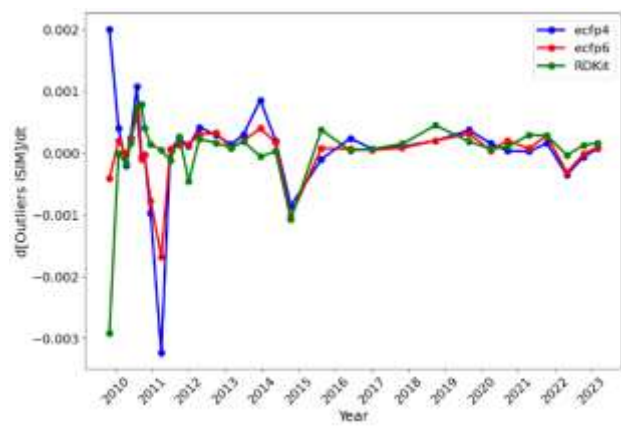

**Figure S13.** iSIM speed respect to size for the A) entire, B) the medoids, and C) outliers of the ChEMBL natural products.

A)

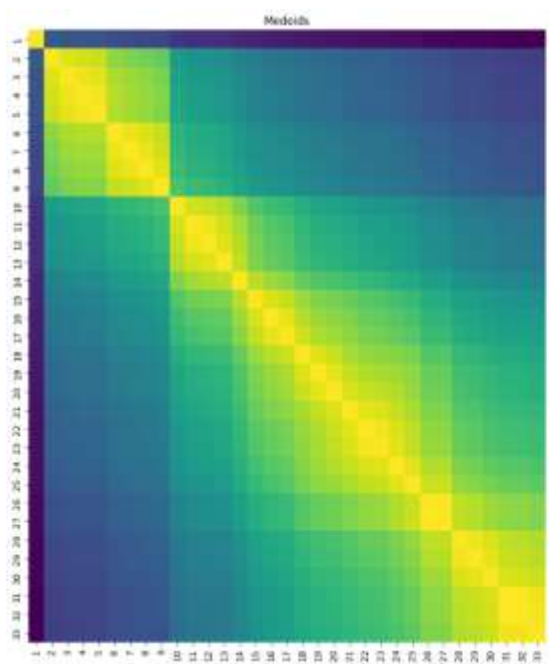

B)

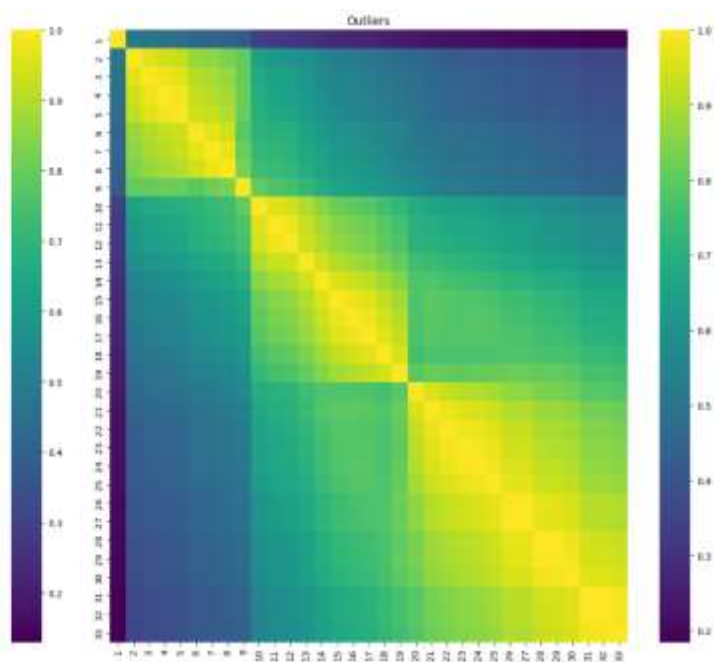

**Figure S14.** Jaccard set similarity values of the medoid (A) and outlier (B) regions of the ChEMBL natural products library releases represented with RDKit fingerprints.

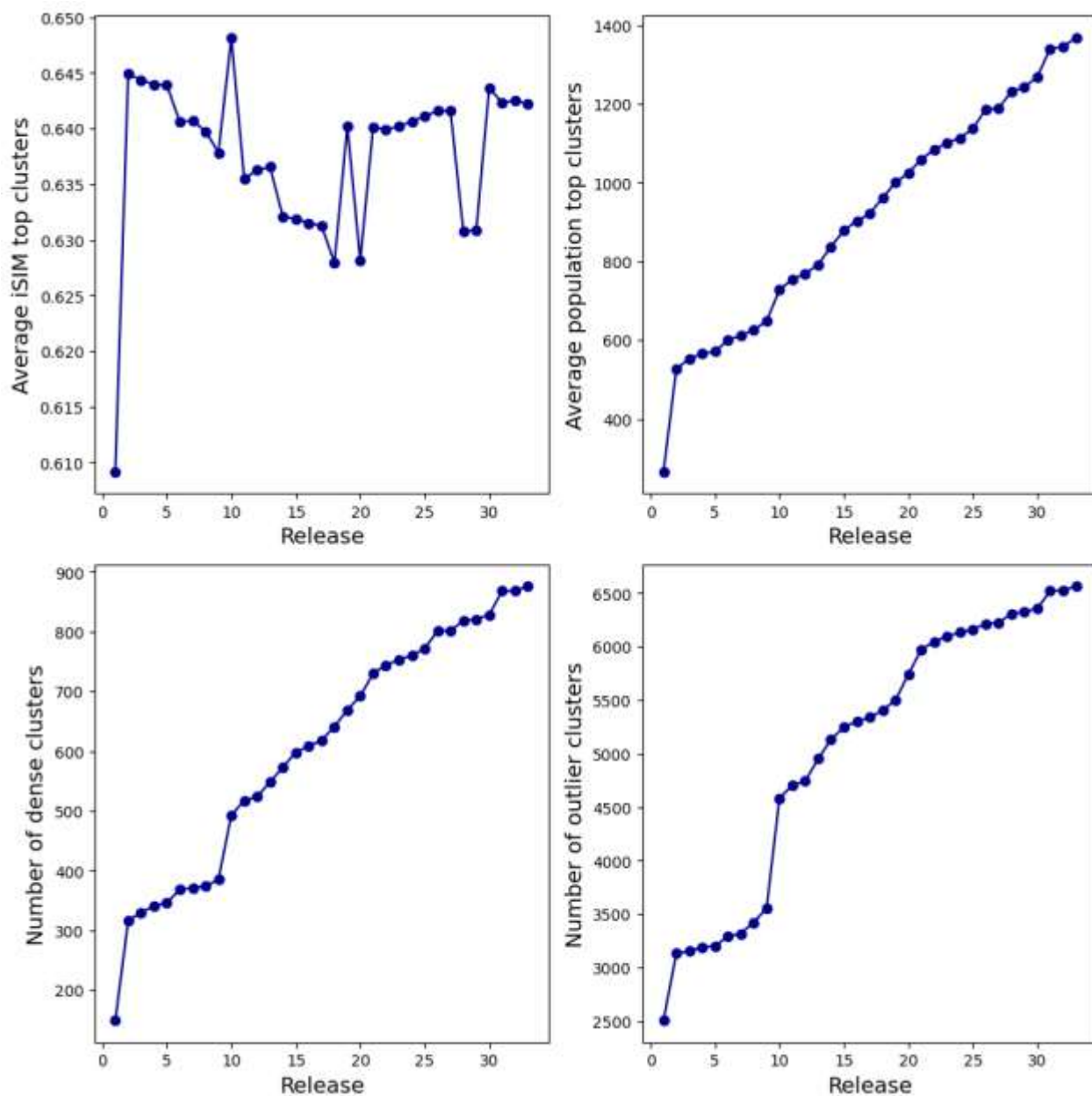

**Figure S15.** (A) Average iSIM of the top 10 most populated clusters (B) Average population of the top 10 most populated clusters (C) Number of dense clusters (D) Number of outliers for the BitBIRCH clustering of the ChEMBL natural products releases over time represented with RDKit fingerprints.

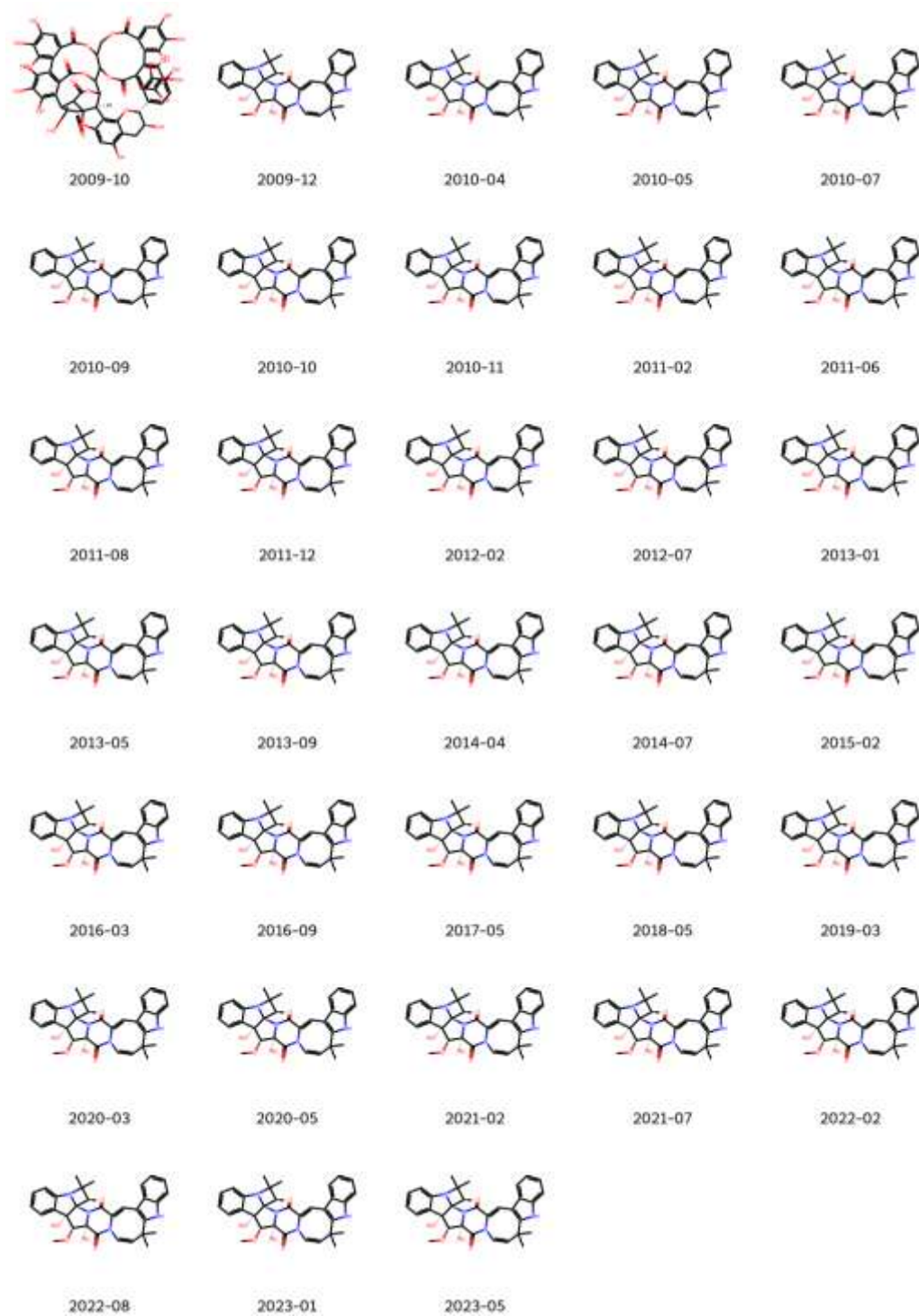

**Figure S16.** Medoids over time for ChEMBL natural products.

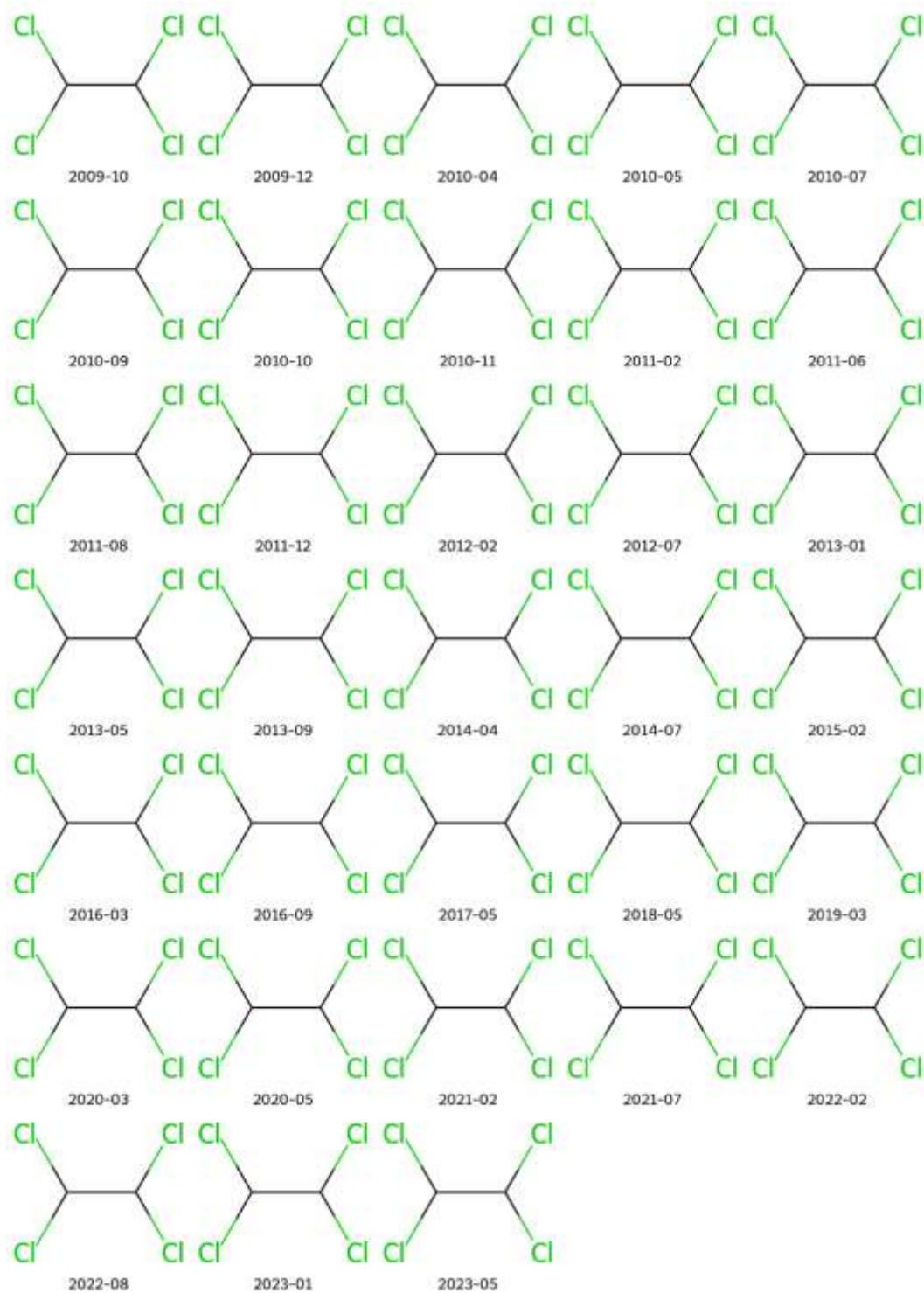

**Figure S17.** Outliers over time for ChEMBL natural products.

- DrugBank

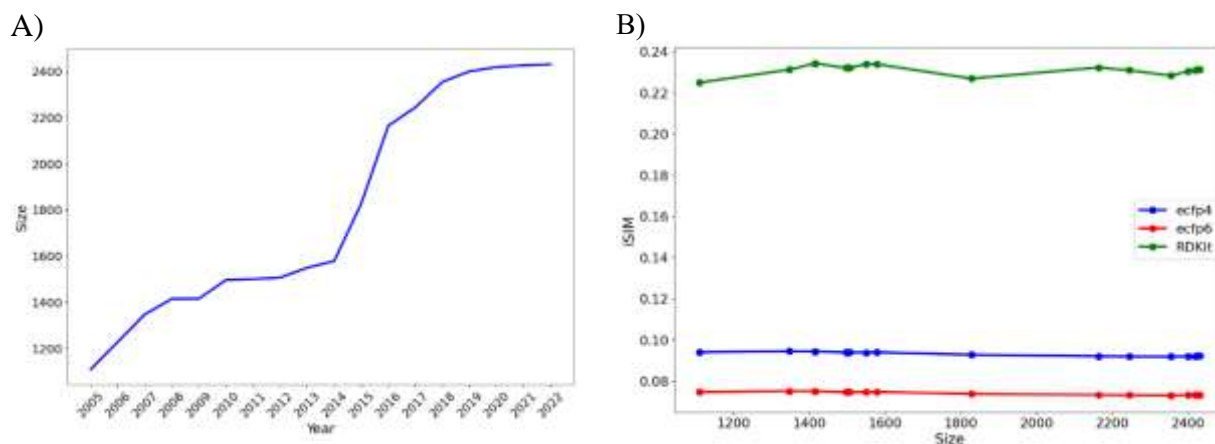

**Figure S18.** (a) Size evolution of DrugBank across releases (2005-2022) over time and (b) the variation of iSIM-Tanimoto with database size for the same releases.

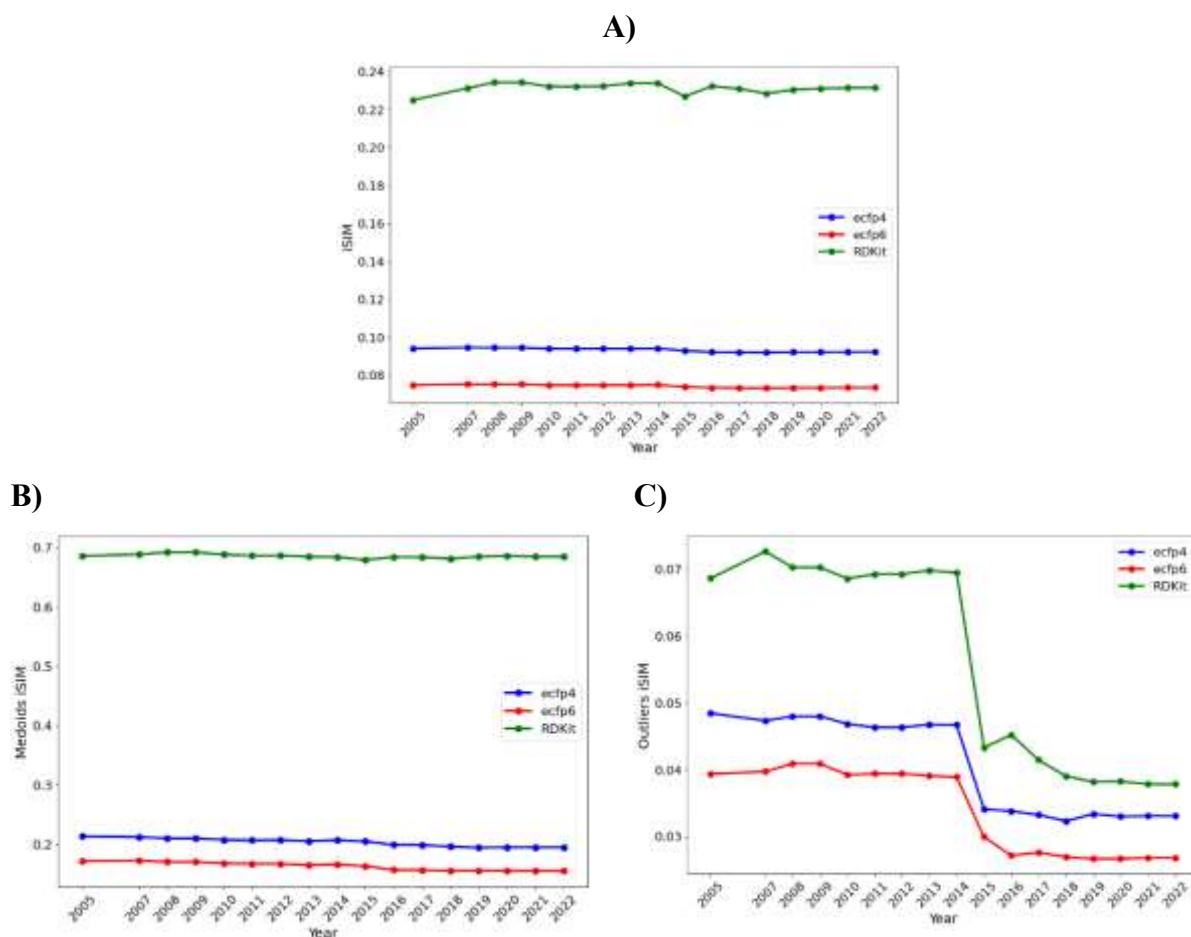

**Figure S19.** Variation of iSIM over time for the (A) DrugBank, (B) medoids, and (C) outliers. Medoids and outliers are considered the 5% of the set with the lowest/highest complementary similarity, respectively.

A)

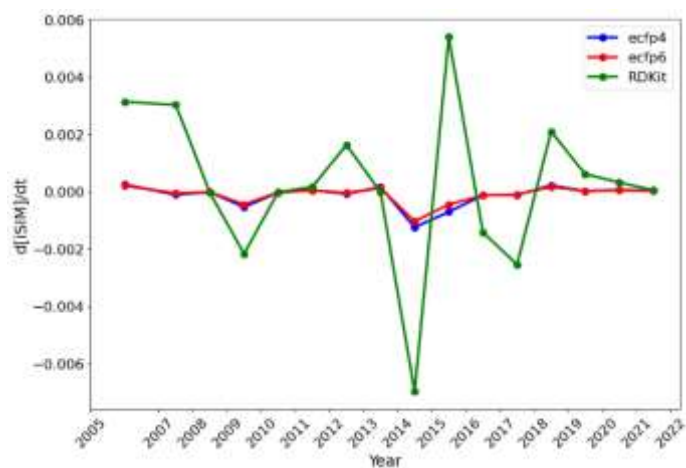

B)

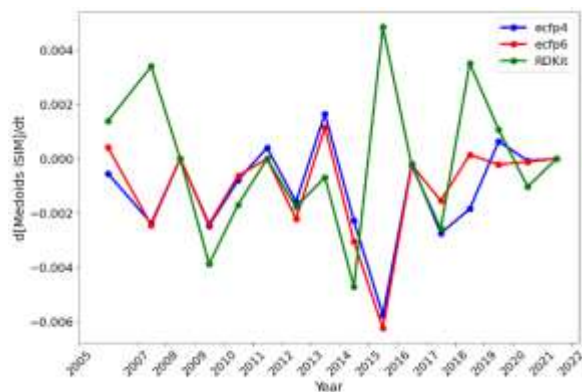

C)

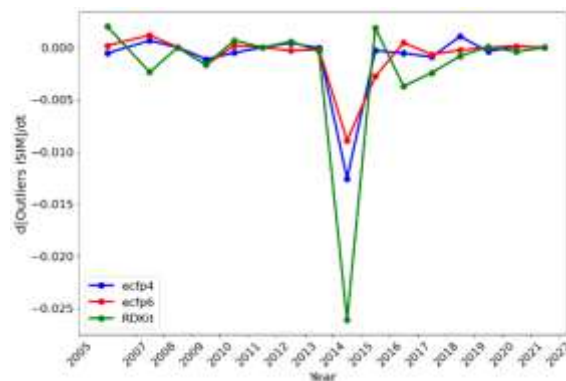

**Figure S20.** Variation of iSIM sped over time for the (A) entire DrugBank, (B) medoids, and (C) outliers. Medoids and outliers are considered the 5% of the set with the lowest/highest complementary similarity, respectively.

A)

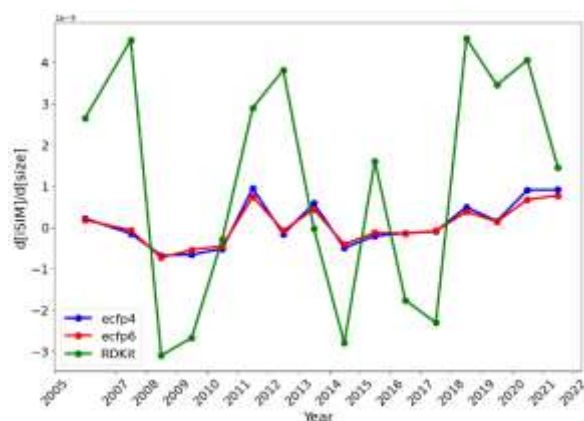

B)

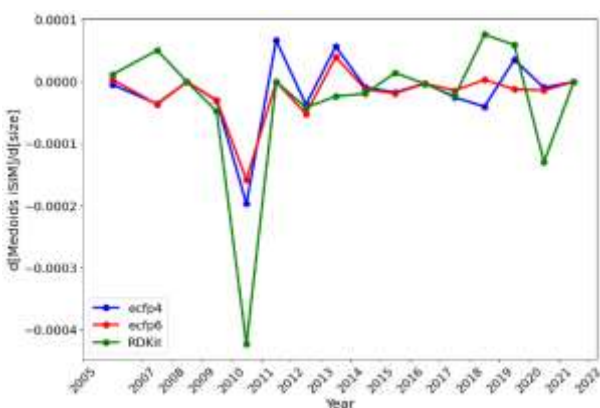

C)

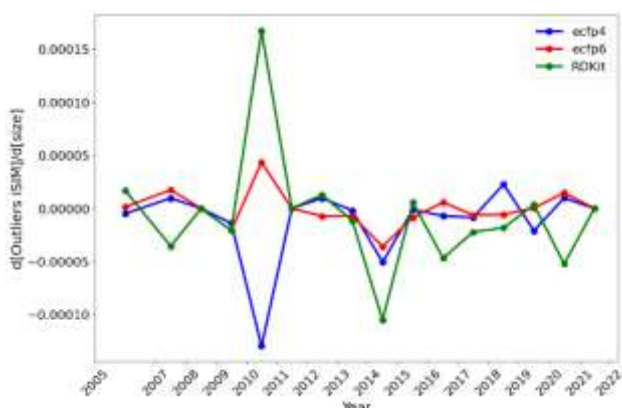

**Figure S21.** Variation of iSIM speed respect to size over time for the (A) entire DrugBank, (B) medoids, and (C) outliers. Medoids and outliers are considered the 5% of the set with the lowest/highest complementary similarity, respectively.

A)

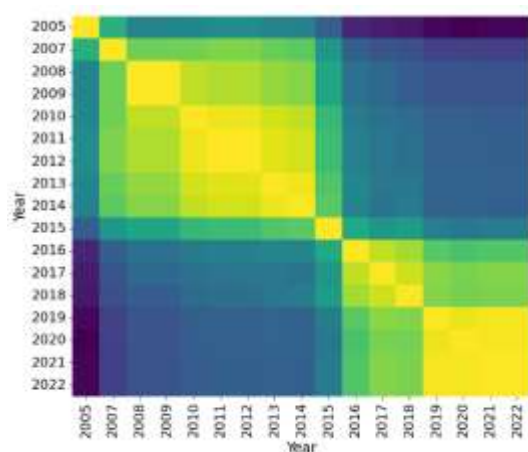

B)

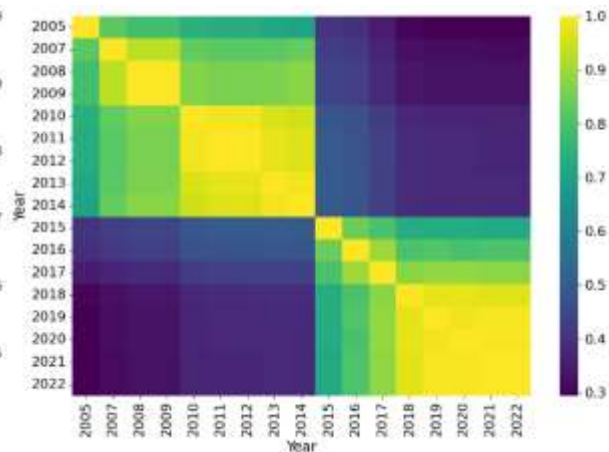

**Figure S22.** Jaccard set similarity values of the medoid (A) and outlier (B) regions of DrugBank library releases represented with RDKit fingerprints.

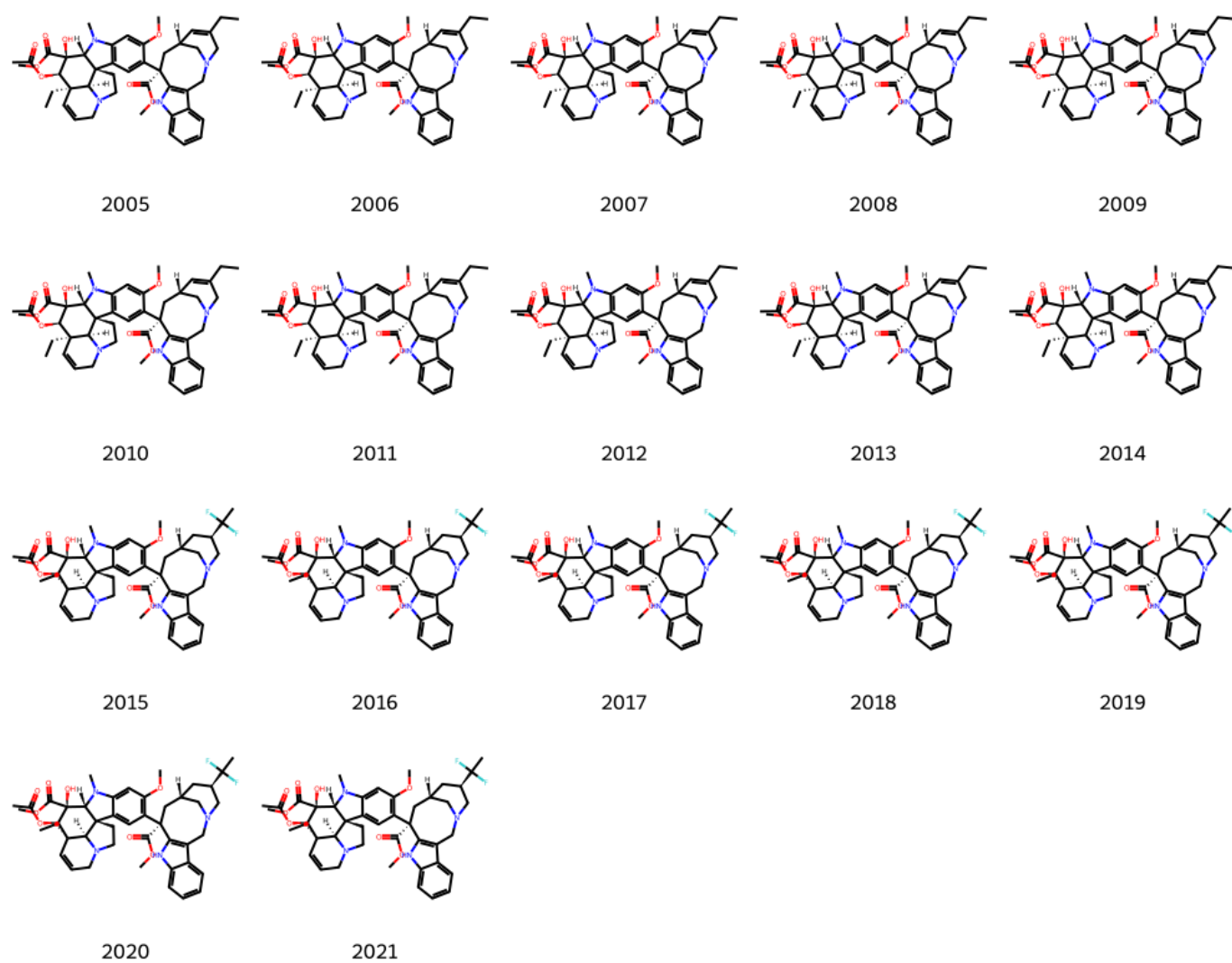

**Figure S23.** Medoids over time for DrugBank.

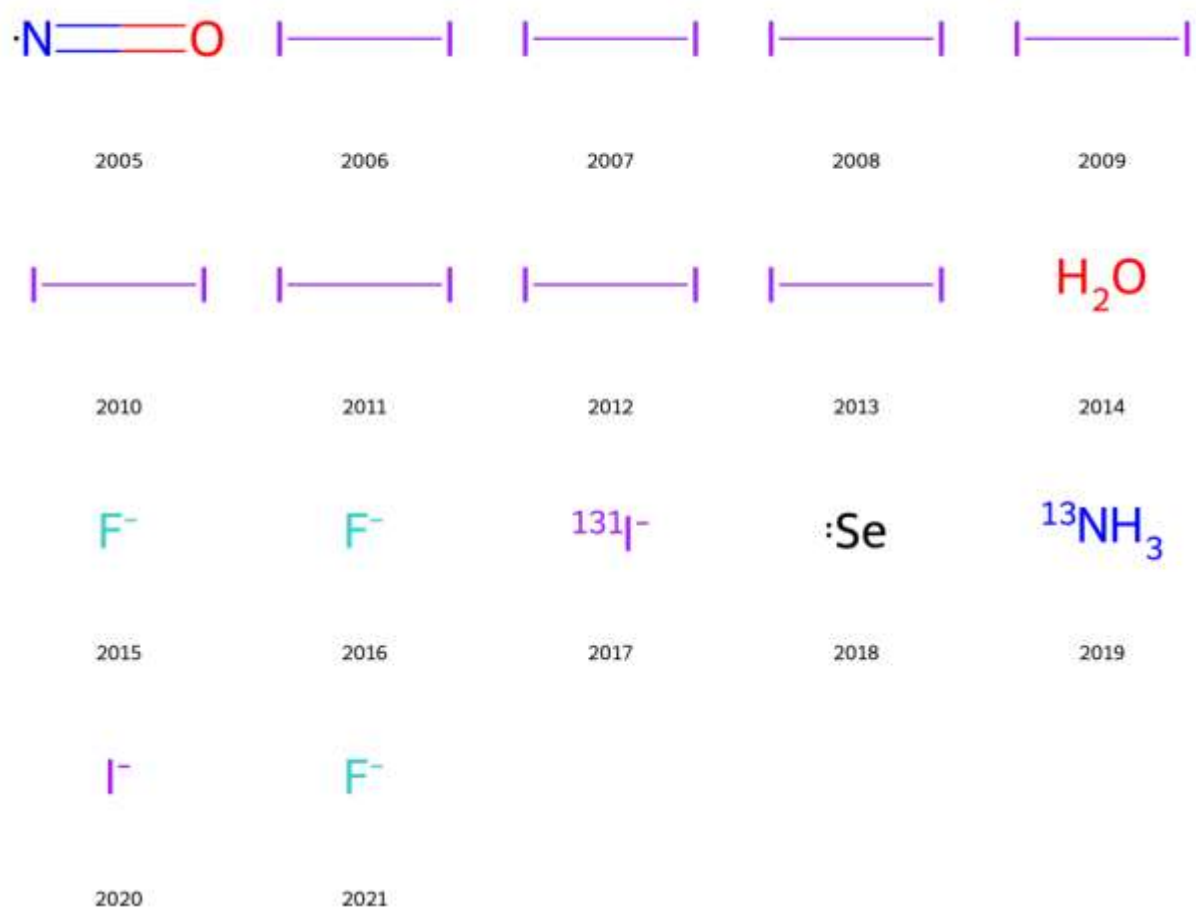

**Figure S24.** Outliers over time for DrugBank.

- *PubChem*

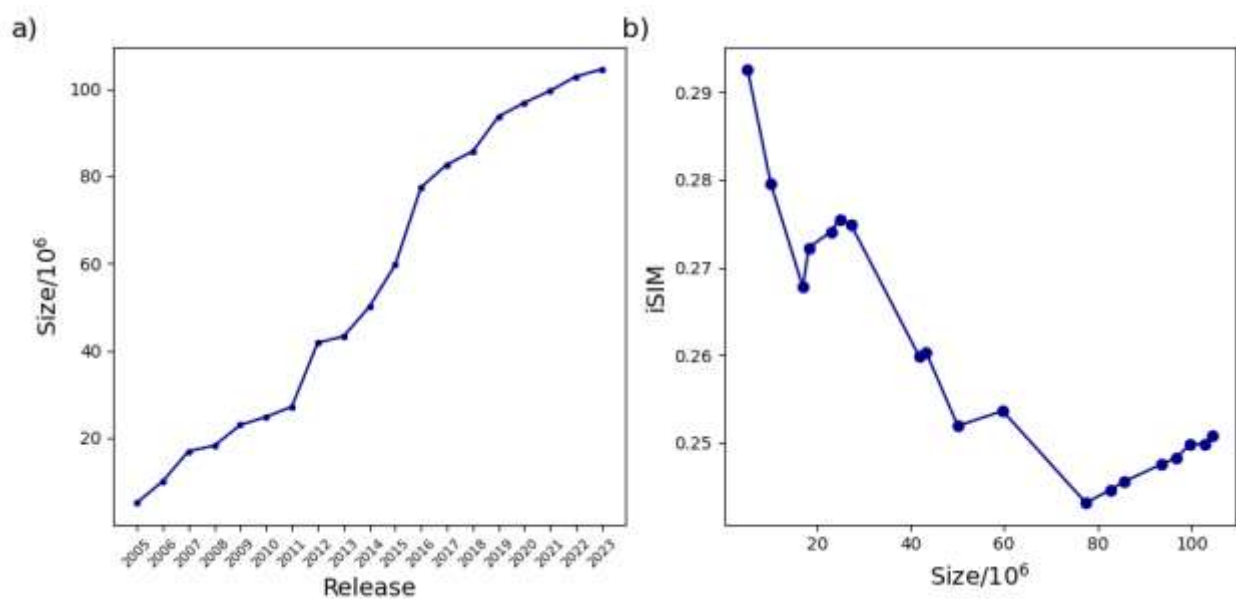

**Figure S25.** (a) Size evolution of the PubChem across releases over time and (b) the variation of iSIM-Tanimoto with database size for the same PubChem releases represented with RDKit fingerprints.

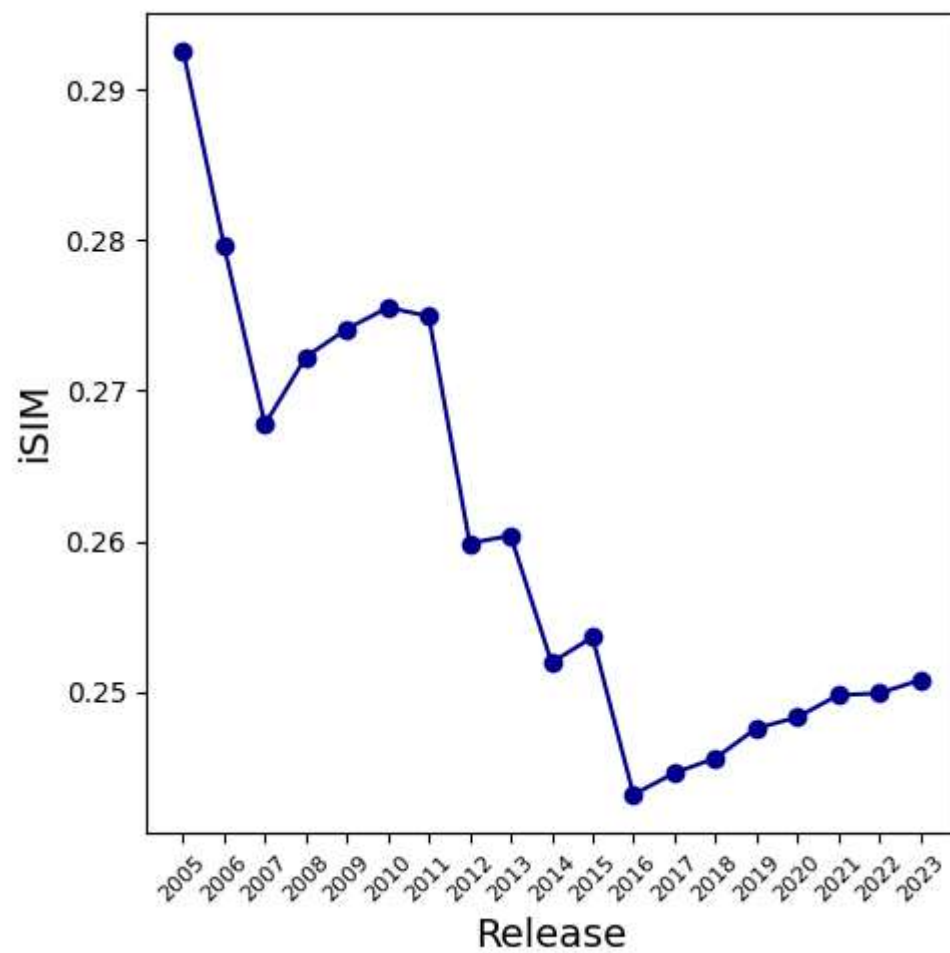

**Figure S26.** Variation of iSIM over time for the PubChem represented with RDKit fingerprints.

A)

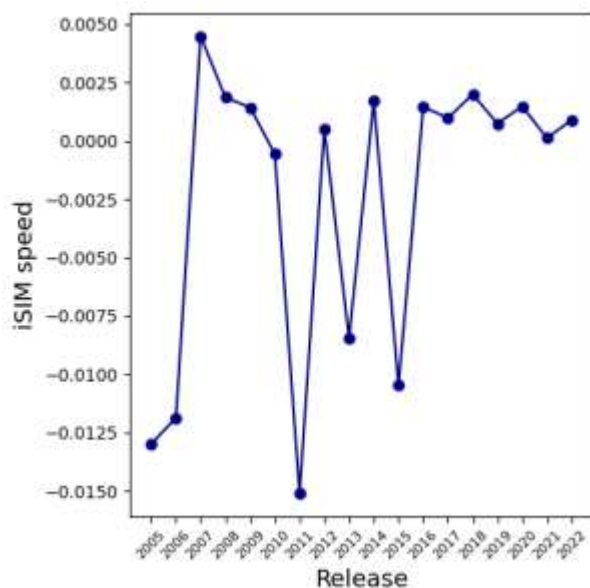

B)

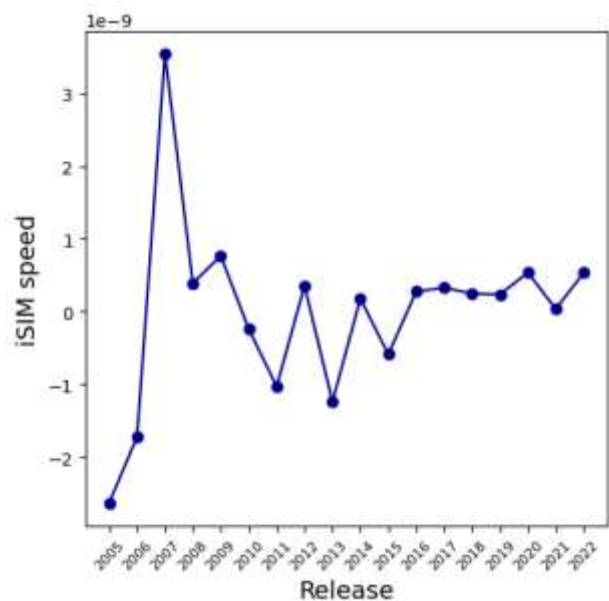

**Figure S27.** iSIM speed respect to A) time and B) size for the PubChem database represented with RDKit fingerprints.

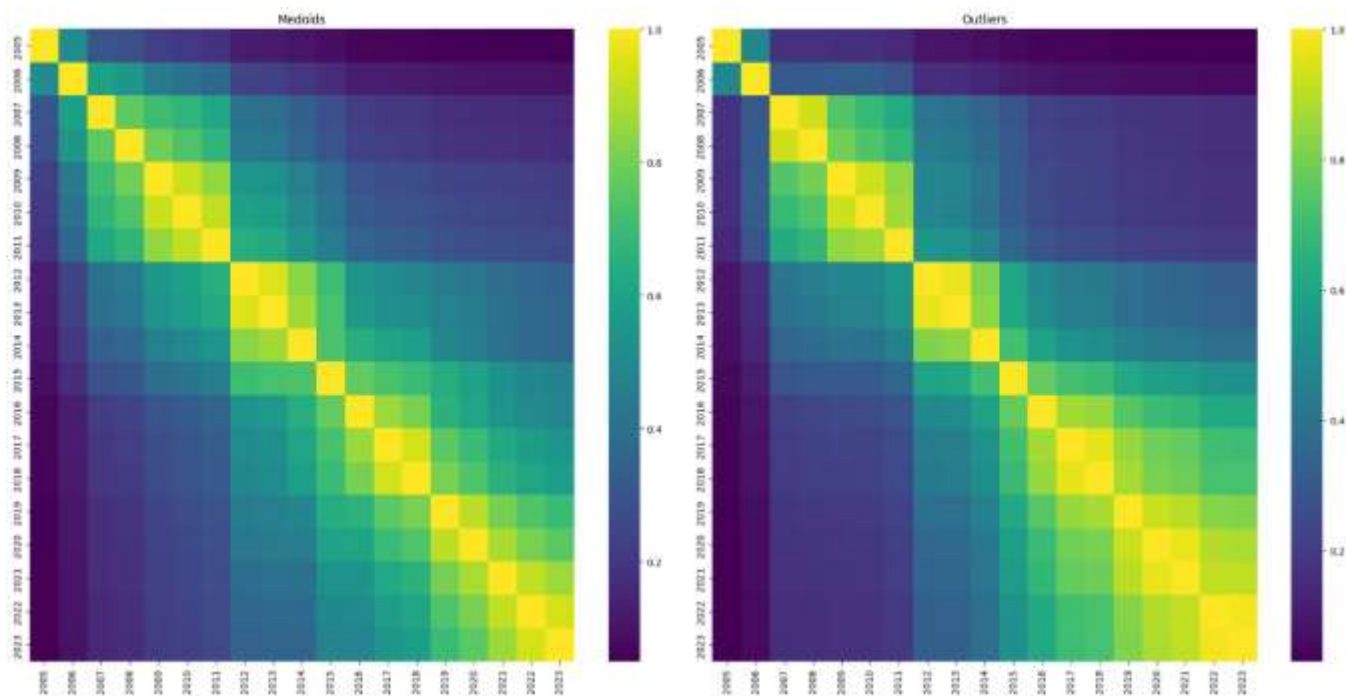

**Figure S28.** Jaccard set similarity values of the medoid (A) and outlier (B) regions of PubChem library releases represented with RDKIT fingerprints.
